# Spaceflight alters host-gut microbiota interactions

**DOI:** 10.1101/2024.01.18.576275

**Authors:** E. Gonzalez, M.D. Lee, B.T. Tierney, N. Lipieta, P. Flores, M. Mishra, N. Beckett, F. Karouia, R. Barker, R.J. Jansen, S.J. Green, S. Weging, J. Broddrick, J. Kelliher, N.K. Singh, D. Bezdan, J. Galazska, N.J.B Brereton

## Abstract

The rodent habitat on the International Space Station has provided crucial insights into the impact of spaceflight on mammals, including observation of symptoms characteristic of liver disease, insulin resistance, osteopenia and myopathy. Although these physiological responses can involve the microbiome when observed on Earth, changes in host-microbiota interactions during spaceflight are still being elucidated. Here, NASA GeneLab multiomic data from the Rodent Research 6 mission are used to determine changes to gut microbiota and murine host colon and liver gene expression after 29 and 56-days of spaceflight. Using hybrid amplicon and whole metagenome sequencing analysis, significant spaceflight-associated alterations to 42 microbiome species were identified. These included relative reductions of bacteria associated with bile acid and butyrate metabolism, such as *Extibacter muris* and *Dysosmobacter welbionis.* Functional prediction suggested over-representation of fatty acid and bile acid metabolism, extracellular matrix interactions, and antibiotic resistance genes within the gut microbiome, while host intestinal and hepatic gene expression described corresponding changes to host bile acid and energy metabolism, and immune suppression from spaceflight. Taken together, these changes imply that interactions at the host-gut microbiome interface contribute to spaceflight pathology and highlight how these interactions might critically influence human health and the feasibility of long-duration spaceflight.

## 1 Introduction

The International Space Exploration Coordination Group, representing 27 of Earth’s space agencies, has outlined a clear target for a crewed mission to Mars in the Global Exploration Roadmap^1,2^, and sustainable long-term lunar exploration as a platform to develop the capabilities necessary to enable this ambitious goal. These guiding objectives have driven development of the imminent commercial low Earth orbit (LEO) destinations and Gateway, and the Artemis mission goal of a permanent lunar surface habitat by the early 2030s^3^. Major challenges associated to longer duration spaceflight and habitation off-Earth are identified in the NASA Moon to Mars Objectives^4^, including the goal to advance understanding of how biology responds to the Moon, Mars, and deep space to support safe human space missions.

Consistently observed spaceflight-associated pathologies, notably disrupted glucose metabolism characterized by insulin resistance and lipid metabolism dysregulation, pose significant risks to astronaut health ^5,6^. Research in tissue culture using the high aspect ratio vessel simulated microgravity model system developed at the NASA Johnson Space Centre characterised increases in pancreatic production of α-TNF, which increased insulin resistance and decreased glucose utilisation in adipocytes^7^. In mice, a reduction of insulin sensitivity has been observed after microgravity simulation using hindlimb unloading^8^. This is reflected in the muscle transcriptome after spaceflight, where insulin receptor signalling is suggestive of disrupted glucose homeostasis^9^.

Similarly, simulated microgravity on human oligodendrocyte^10^ and mesenchymal stem^11^ cell cultures increases production of fatty acids and complex lipids. In the nematode *Caenorhabditis elegans*, the intestinal lipid metabolic sensors SBP-1 and MDT-15 respond to simulated microgravity, with RNAi knockdown of *sbp-1* and *mdt-15* reducing lipid toxicity^12^. Spaceflight metabolic studies from the Bion space program (Kosmos 605, 690, 782, 936 and 1887 (1973-87))^13–15^ characterised rats as hyperlipidemic, with spaceflight inducing elevated serum or hepatic fatty acids, and substantial increases in cholesterol (67%). Similar lipid dysregulation, suggestive of non-alcoholic fatty liver disease (NFALD), has been a consistently observed mammalian response to spaceflight alongside aligned disruption of insulin metabolism and glucose homeostasis^16–20^. These observations in mice and humans on the ISS include widespread changes in the hepatic proteins which drive lipid metabolism, significant increases in steatosis, cholesterol and low-density lipids and reduced high-density lipids.

The immune system can be compromised by spaceflight, both in space and after return to Earth. Despite quarantine before flight, infection with influenza and *Pseudomonas aeruginosa* have been observed in astronauts^21^. Up to 50% of astronauts also exhibit immunodeficiency upon returning to Earth^22^, leaving them vulnerable to infection. This dysregulation manifests through decreased T cell and B cell abundance^23^, and impaired natural killer cell and macrophage function^24,25^. The underlying cause of these changes are thought to be driven by microgravity, isolation, and stress associated with spaceflight^26^, as well as shifts in the gut microbiome^27^. On Earth, comparable changes in muscle integrity, glucose homeostasis, lipid metabolism, immune and psychophysiological function have been associated to gut microbiota^28–32^. Similarly, unique built environment surface microbiology arises from long-duration confinement, reshaping the bidirectional exchanges between usually diverse environmental microbial ecosystems and the gut microbiome to promote opportunistic pathogenicity^33–37^.

Given the potential involvement of gut microbiota in spaceflight pathology, and their essential role in mediating healthy human metabolic function on Earth, there has been increasing research into gut microbiome dynamics associated with spaceflight. Using 16S ribosomal RNA gene (16S rRNA) amplicon sequencing, Jiang et al^38^ identified significant changes in the relative abundance of 16 OTUs in the gut microbiome of mice in Rodent Research (RR) 1 (RR-1) mission, some of which were annotated as within the genera *Staphylococcus* and *Tyzzerella*, and were lower in mice after spaceflight compared to ground controls. More recently, Bedree et al.^39^ explored the gut microbiome of mice flown in the RR-5 mission using 16S rRNA amplicon sequencing and whole metagenome sequencing (WMS). Amplicon analysis identified 14 ASVs as different in relative abundance (p < 0.05) between spaceflight (ISS) and ground controls after 9 weeks of spaceflight, including increases in the genera *Clostridium*, *Romboutsia*, *Ruminiclostridium*, and *Shuttleworthia*, and decreases in *Hungatella*, while WMS identified significant enrichment of *Dorea* sp. and the species *Lactobacillus murinus*.

In this study, species-resolved 16S rRNA amplicon sequencing and *de novo* co-assembled WMS were employed to capture metagenomic changes in the murine gut microbiome associated with spaceflight across multiple samples as part of the RR-6 mission (Fig 1AB). Intestinal and hepatic transcriptomics were then used to assess the associated gene expression response of mice to spaceflight.

**Figure 1.**
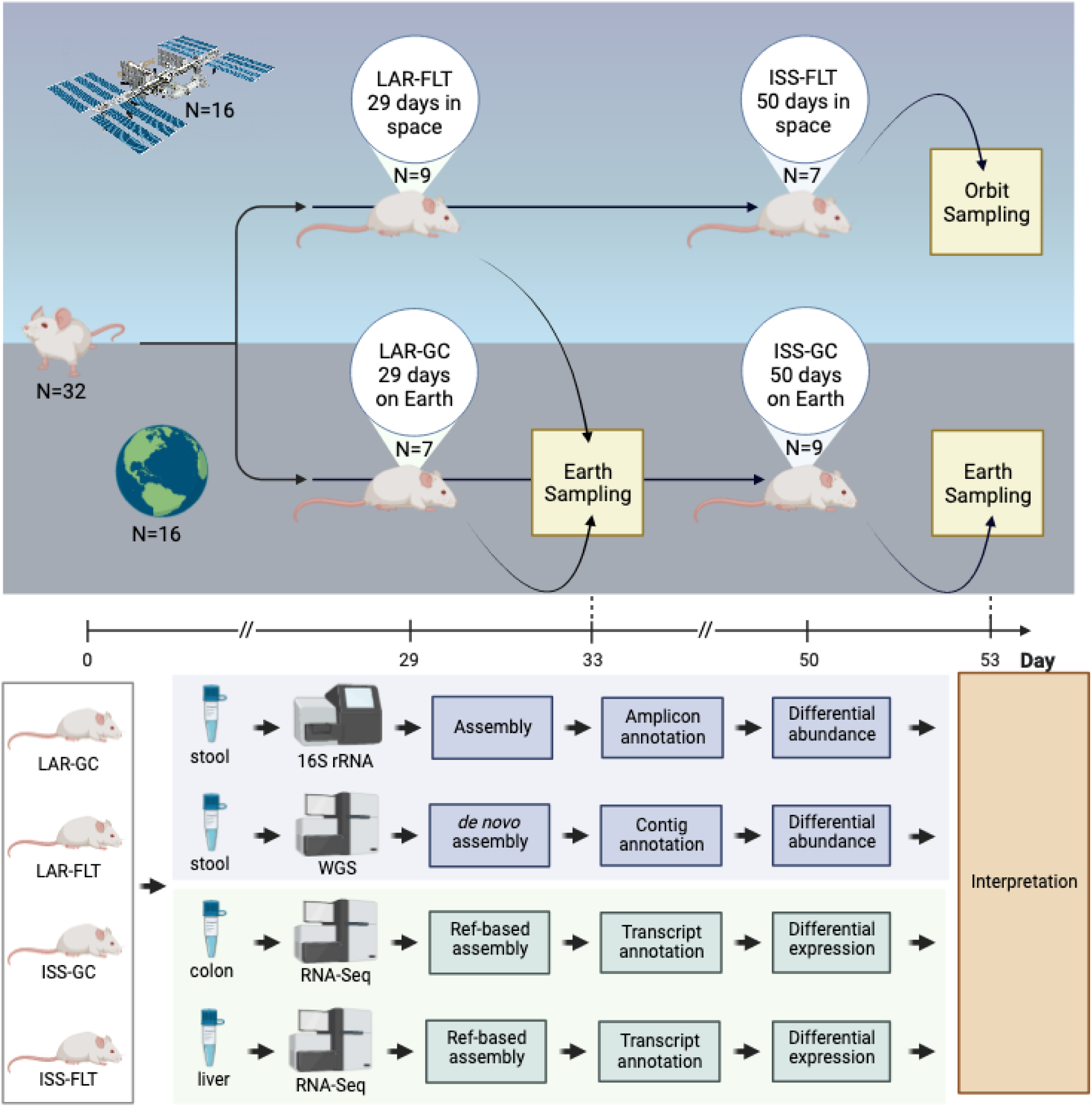
Experimental design. Analysis design of data from the Rodent Research 6 mission and multiomic data analysis strategy

## 2 Results and discussion

### 2.1 Spaceflight increased total body weight

Although reduced muscle mass and bone density in astronauts and mice during spaceflight are commonly observed^40,41^, in this study, total mouse body weight was trending to increase after 29 days of spaceflight in Live Animal Return mice (FLT_LAR; n= 9) and significantly increased after 56 days of spaceflight in ISS mice (FLT_ISS; n= 7, p<0.05). Carcass mass did not differ significantly (t-test, p>0.05) between Ground Control Live Animal Return (GC_LAR; n= 7) and FLT_LAR mice, which weighed 28.9 g^-1^ (±1.6) and 30.1 g^-1^ (±1.4), respectively, but did significantly (t-test, p<0.05) increase from 28.4 g^-1^ (±1.4) in Ground Control ISS (GC_ISS; n = 9) mice to 32.9 g^-1^ (±1.0) in FLT_ISS mice (Supplementary document 1). Suzuki et al.^42^ observed similar increases in the mouse habitat unit 3 mission and attributed these changes to substantial increases in both white and brown adipose tissue, and large increases in total plasma cholesterol and triglyceride levels. While healthy adipose cells play an important role in maintaining insulin sensitivity, dysregulated adipose can lead to production of pro-inflammatory and insulin-antagonistic molecules^43,44^.

### 2.2 Spaceflight alters murine gut microbiota

Insulin resistance and lipid accumulation are common spaceflight phenotypes^45^ which are influenced by short chain fatty acids (SCFAs) and can be improved through butyrate dietary interventions in ground-based murine studies^46,47^. As butyrate and other SCFAs are predominantly produced by bacteria within the gut^48^, alterations in RR6 gut microbiome composition were explored. Characterisation of microbiota used 16S rRNA amplicon sequencing as well as WMS sequencing from faecal samples collected from GC_LAR and FLT_LAR (after 29 days of spaceflight) as well as GC_ISS and FLT_ISS mice (after 56 days of spaceflight).

Sequencing of amplicon libraries generated 2,146,311 sequences after quality control, with an average of 77,015 ± 853 per sample (Fig 2A-F; Supplementary file 1). A total of 133 exact sequence variants (ESVs) were inferred across all samples; these ESVs accounted for 1,959,722 (91.32%) of reads. Thirty-five ESVs were annotated as putative bacterial species, 12 at genera level and 11 at family level, with an average of >99.9% nucleotide identity, while 74 ESVs were dissimilar to any well-characterised taxa (<99% nucleotide identity; 14 of which were flagged as putative chimeras). Most counts (73.1%) were captured by ESVs annotated as putative species, 24 of these could be assigned to a single species and 11 to multiple species which share identical rRNA gene sequences at the V4 region of the 16S rRNA gene. Bacterial species putatively identified across all mice using amplicons belonged to the phyla Firmicutes (28), Proteobacteria (4), Actinobacteria (1), Deferribacteres (1) and Bacteroidetes (1). *Parabacteroides goldsteinii* had the highest relative abundance throughout the samples and accounted for 53.7% of sequence counts.

**Figure 2.**
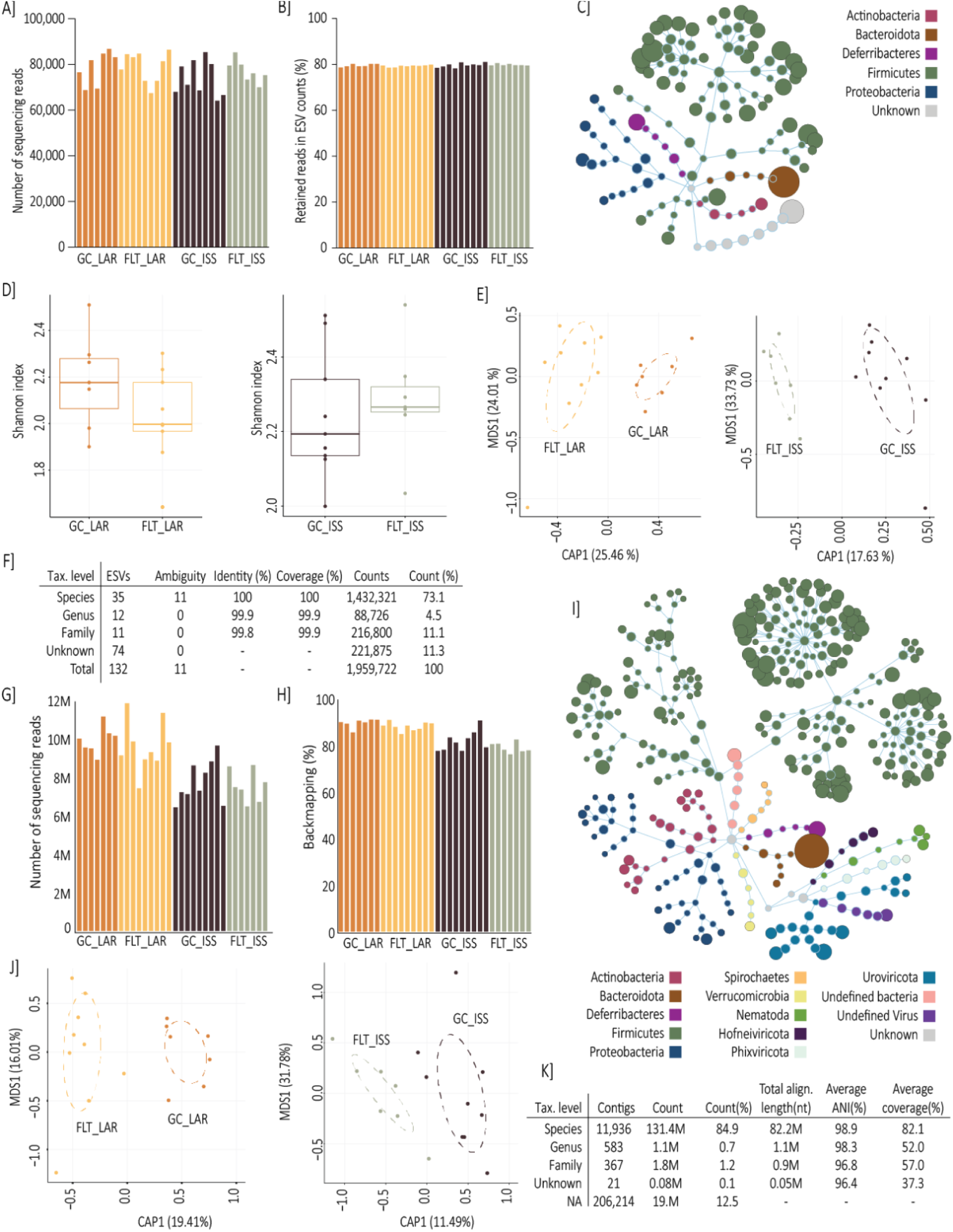
Ground control, live animal return and ISS murine gut microbiome capture. Amplicon library sizes for A) GC_LAR and ISS_LAR, and B) GC_ISS and FLT_ISS. C) Flower diagram illustrating amplicon phyla diversity. D) Amplicon alpha diversity. E) Amplicon Canonical Correspondence Analysis (CCA). F) Amplicon annotation and count distribution summary statistics. G) and H) WGS library sizes. I) Flower diagram illustrating WGS phyla diversity. J) WGS CCA. K) WGS annotation and count distribution summary statistics. Extended details in Supplementary files 1-3.

WMS of faecal DNA generated 277,294,078 reads after quality control, with an average of 8,665,440±1,423,541 per sample (Fig 2G-K; Supplementary file 2). Co-assembly across all 32 mice generated 219,259 contigs and back-mapping captured 85.1% of raw counts after mouse filtering (154,692,394 counts). After sparsity (>90% counts in a single sample) and minimum sample (<3) occurrence filtering to allow statistical detection of spaceflight associated differences across biological replicates, 45,890 contigs remained which captured 154,234,982 counts. These contigs were provisionally annotated (>90% nucleotide identity) as putative bacteria (27.6%), viruses (0.8%) and metazoa (0.1%), or were unknown (71.5%), and included 11,936 contigs which could be provisionally annotated from 162 species and shared an average of 98.93% average nucleotide identity (ANI) with known species or strains (and capturing 84.96% of total counts) (Fig 2K; Supplementary file 3). Grouping WMS contigs by species annotation and filtering for high confidence (>97% ANI, and >2000nt total length) identified 79 species of bacteria representing 66 species within Firmicutes, and 6 Actinobacteria, 4 Proteobacteria, 1 Bacteroidetes, 1 Spirochaetes and 1 Deferribacteres, as well as the helminth, *Trichinella nativa*. These species groups ranged from 1 to 2056 contigs, with an average total alignment length of >950,000 nt and an ANI of 98.8% (Supplementary file 3). Prevalent taxa included *P.goldstienii* representing 37%, *Enterocloster clostridioformis* representing 5% and all other species representing below 1% of relative abundance across all mice, including non-bacterial species such as *Trichinella nativa,* representing 0.04% of relative abundance and present in all mice. *P. goldsteinii* is a ubiquitous commensal gut microbiome inhabitant in mice, and the species includes strains that play a role in reducing intestinal inflammation and maintaining intestinal epithelial integrity^49,50^.

Microbial alpha diversity indices did not significantly differ between mice groups using 16S rRNA amplicon sequencing or WMS data (Fig 2D; Supplementary file 1). Canonical Correspondence Analysis showed both FLT_ISS and GC_ISS as well as FLT_LAR and GC_LAR samples segregated by group, and the first axes explained 17.63% and 25.46% of variation using 16S rRNA gene amplification and 19.41% and 11.49% of variation using WMS, respectively. ANOVA-like permutation tests confirmed significant variation between groups under constraint for both amplicon and WMS data (Fig 2EJ; p < 0.05; Supplementary file), suggesting spaceflight influenced gut microbiota in both comparisons regardless of the metagenomic approach taken.

#### 2.2.1 Significant spaceflight changes in the microbiome community are associated with short-chain fatty acid metabolism, bile acid conversion and pathogenicity

Differential abundance analysis of amplicon data identified 45 ESVs that were significantly different in relative abundance between spaceflight and ground control mice, including 34 ESVs between GC_LAR and FLT_LAR and 18 ESVs between GC_ISS and FLT_ISS (Fig 3A and B). Although there were divergent changes in the relative abundance of *H. xylanolytica,* the common significant enrichment of *E. muris* and *D. welbionis* in mice after 29 and 56 days of spaceflight, compared to separate matched control groups, suggests spaceflight had some common influence of gut microbiota which persisted over the 29-56 days onboard the ISS as well as some distinct effects over time.

**Figure 3.**
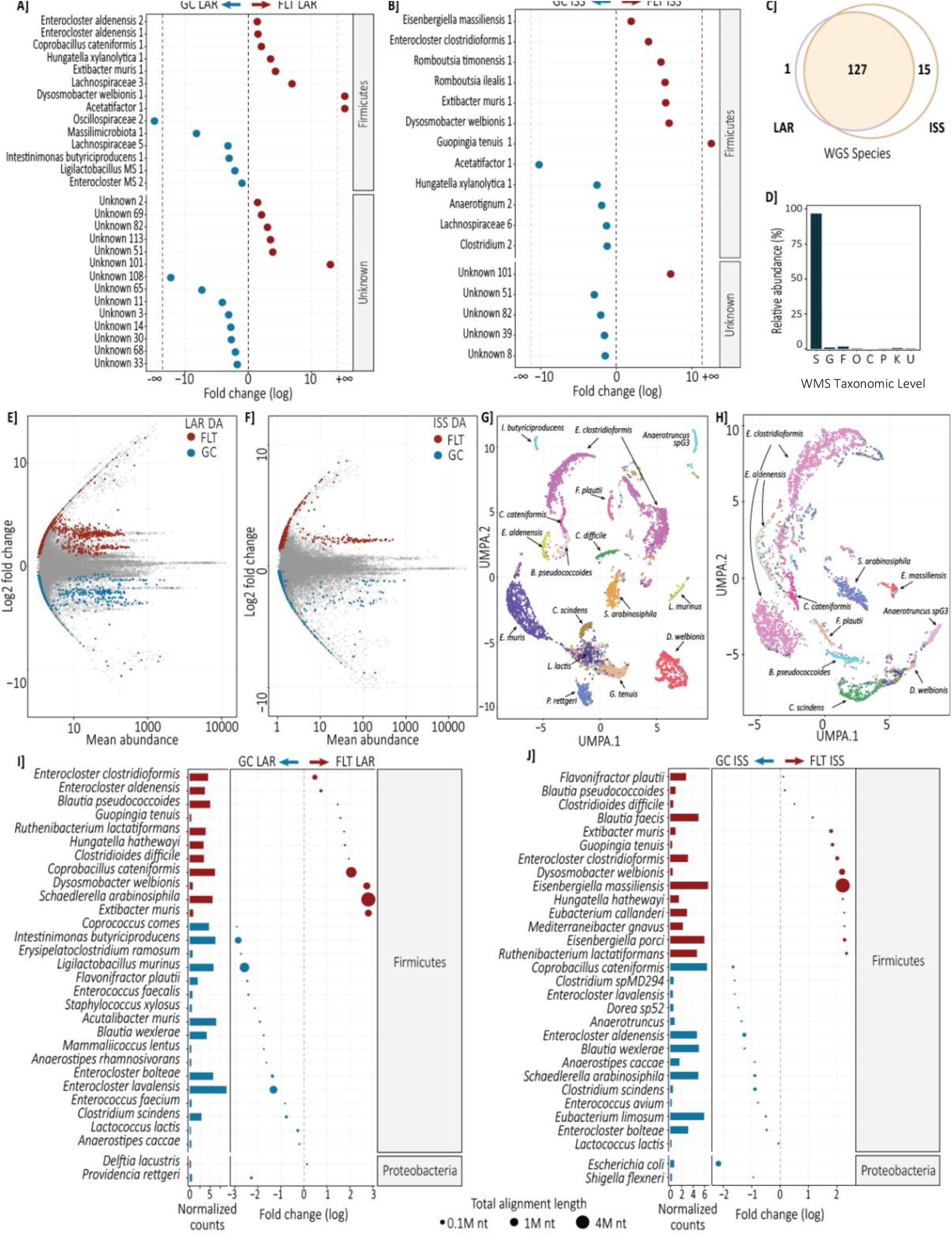
Spaceflight-assoicated significant micorbiome alterations. Significantly differentially abundant (DESeq2) ESVs between A) GC_LAR and ISS_LAR, and B) GC_ISS and FLT_ISS. Fold change (FC log2) in relative abundance. +/− INF (demarcated by the dashed red line) indicates ‘infinite’ fold change, where an ESV had detectable counts in samples from only one condition (structural zero). C) Comparison of WGS detected species between LAR and ISS samples. D) Distribution of counts across WGS taxonomy. E) and F) Contig WGS MA plots with significantly differentially abundant (DESeq2) highlighted. G) and H) UMAP diagrams used to visualise contig clustering of selected species and J) Significantly differentially abundant (DESeq2) species detected with WGS, node size illustrates contig number. Extended details in Supplementary files 1-3. Amplicon sequencing: Species enriched in FLT_LAR mice compared to controls included *Coprobacillus cateniformis, Dysosmobacter welbionis*, *Enterocloster aldenensis*, *Extibacter muris* and *Hungatella xylanolytica*, while depleted species included *Intestinimonas butyriciproducens* and ESVs ambiguous to multiple *Enterocloster* species (including *E.lavalensis*) and *Ligilactobacillus* species (including *L.murinus*). Species enriched in FLT_ISS mice included *D.welbionis*, *Eisenbergiella massiliensis, Enterocloster clostridioformis, E.muris*, *Guopingia tenuis*, *Romboutsia ilealis* and *Romboutsia timonensis*, while depleted species included *H.xylanolytica*. WMS: Microbiome species significantly enriched in after 29 days of spaceflight comprised 11 Firmicutes, including *Blautia pseudococcoides*, *Clostridioides difficile*, *C.cateniformis*, *D.welbionis, E.aldenensis*, *E.clostridioformis*, *E.muris*, *G.tenuis*, *Hungatella hathewayi*, *Ruthenibacterium lactatiformans, Schaedlerella arabinosiphila* and the proteobacteria *Delftia lacustris*. Significantly depleted species included 18 firmicutes, including *Acutalibacter muris, Anaerostipes caccae, Blautia wexlerae, Clostridium scindens, Enterococcus faecalis, Ligilactobacillus murinus, Enterocloster bolteae, E.lavalensis, Flavonifractor plautii*, *I.butyriciproducens*, *Lactococcus lactis and Staphylococcus xylosus*, and the Proteobacteria *Providencia rettgeri*. These findings agreed with significant differential abundance of *C.cateniformis, D.welbionis, E.aldenensis*, *E.clostridioformis*, *E.muris* and *I.butyriciproducens* inferred from 16S rRNA amplicon analysis and resolved species ambiguity for *E.bolteae, E.lavalensis* and *L.murinus*. Microbiome species which were significantly enriched in after 56 days of spaceflight comprised 14 Firmicutes, including *B.pseudococcoides, C.difficile, D.welbionis, E.clostridioformis, Eisenbergiella massiliensis, E.muris, F.plautii, G.tenuis, H.hathewayi* and *R.lactatiformans.* Significantly depleted species included 17 Firmicutes, including *A.muris, Anaerostipes caccae*, *B.wexlerae*, *C.scindens*, *C.cateniformis*, *E.aldenensis*, *E.bolteae*, *E.lavalensis*, *L.lactis, S.arabinosiphila* and as well as the Proteobacteria *Escherichia coli* and *Shigella flexneri*.

Differential abundance analysis of WMS identified 13,996 contigs that were significantly different (DESeq2 FDR < 0.1) in relative abundance between spaceflight and ground control mice, including 11,087 between GC_LAR and FLT_LAR, and 3,997 between GC_ISS and FLT_ISS (Fig 3C-H). From these, 30 putative species (99.0% ANI; Supplementary file 3) identified as significantly differentially abundant between GC_LAR and FLT_LAR (Fig 3I) with an average total length of 433,429 nt per species, while 30 species (98.9% ANI) significantly differed between GC_ISS and FLT_ISS (Fig 3J) with an average total length of 191,542nt.

*D. welbionis* was recently characterised by Roy et al^51,52^ as a butyrate producer likely present in the gut of most humans and was negatively correlated with BMI in obese individuals with metabolic syndrome. The same team used murine supplementation experiments to illustrate that *D. welbionis* could partially counteract insulin resistance, adipose tissue hypertrophy and inflammation as well as suggest a potential association with mitochondrial content and activity in adipose tissue after high fat diet induction of obesity^52^. In mice, changes in microbially produced butyrate are also know to directly influence expression of hepatic circadian clock regulating genes, such as *Per2* and *Bmal1*, in a bidirectional interaction which can disrupt host metabolism^53^. Enrichment of *D. welbionis* in both groups of spaceflight mice (fig 3ABIJ) compared to their respective ground controls here is therefore noteworthy given the high lipid accumulation, liver and mitochondrial dysfunction phenotype repeatedly observed in rodent research missions and astronauts^17,18,45^. Whether the relative increase of this species might be counteracting or contributing towards spaceflight pathology is unclear and merits further study. Conversely, other butyrate producers, such as *Intestinimonas butyriciproducens*^54^, were depleted after 29 days of spaceflight.

*L. murinus* and *A. muris* were depleted in mice during spaceflight (fig 3AI), as well as some *Enterocloster* species after 29 days of spaceflight, which can have high expression of bile salt hydrolases (BSHs), able to deconjugate bile salts into less toxic bile acids, and can promote microbially mediated 7α-dehydroxylation of host primary bile acids into secondary bile acids^55–59^. Conversion of the major human primary bile acids in humans, cholic acid (CA) and chenodeoxycholic acid (CDCA), to the secondary BAs deoxycholic acid (DCA) and lithocholic acid (LCA), is mediated by a limited number of closely related clostridia containing the bile acid inducible (bai) operon, such as *Clostridium scindens*^60,61^ which was significantly reduced in abundance after spaceflight (Fig 3IJ). The major murine primary bile acids also include α- and β-muricholic acid (αMCA and βMCA), which are transformed by 7α-dehydroxylation to murideoxycholic acid (MDCA).

*E. muris*, which significantly increased in both spaceflight groups of mice compared to ground controls (Fig 3ABIJ), has been recently characterised as 7α-dehydroxylating in mice^62^, containing the bile acid inducible operons BaiBCDEFGI and BaiJKL, and BaiA, homologous to *Clostridium scindens*. The Bai operon enables *E. muris* and *C. scindens* to increase concentrations of 7α-dehydroxylated secondary BAs that alter the host bile acid pool and act as ligands to bile acid receptors to influence host inflammation, glucose and lipid metabolism^62–66^. For example, bile sensor farnesoid-X-receptor (FXR) modulates enterohepatic recirculation and host cholesterol metabolism through bile acid regulation of *cyp71A*^66^. Similarly, secondary bile acids such as DCA and LCA are potent agonists of the bile acid receptor TGR5, which controls glucose homeostasis in adipose tissue and muscle by altering intestinal cell release of the insulin secretion regulator glucagon-like peptide-1 (GLP-1)^67–69^. Liver production of α-MCA and β-MCA (in mice) is mediated by *cyp2c70* genes^70,71^ but 7α/β-dehydroxylation mediated by microbes such as *E.muris* can modify MCAs after epimerization into HDCA^72^, and critically regulate lipid metabolism^73,74^. Interestingly, *E. clostridioformis*, significantly higher in relative abundance after both 29 and 56 days of spaceflight (fig 3), is reported as increasing in the presence of *E. muris*^62^ and harbours 7α/β hydroxysteroid dehydrogenases (HSDH)^59,75^, which can also transform primary and secondary BAs into oxo-bile acids^64^.

Immune suppression has previously been described as a response to spacefight^76^ and could result as bile acid dysregulation^77,78^. So increased relative abundance of *C. difficile* after both 29 and 56 days of spaceflight is of potential concern if toxigenic.

#### 2.2.2 Changes in metagenome functional prediction

Metagenomic functional prediction identified 4,583,759 genes in the co-assembly generated from all 32 mice, 392,631 of which were annotated by Kegg database (Supplementary file 4). Thus, a high proportion (91.4%) of genes, including differential abundant genes, remain unannotated. Kegg annotated genes included the pathogenicity locus (including *tcdAB*) from *C. difficile*, suggesting significant enrichment of the species after 29 and 56 days (fig 3IJ) could include a toxigenic strain. Bile acid metabolism genes were identified, including 17 bile salt hydrolases, including that from spaceflight enriched *A. muris* (*Cbh*), and 57 non-redundant Bai genes, including from *C. scindens* (*BaiABCDEFGI*) and *I. butyriciproducens* (*BaiA* and *BaiCD*), species which were significantly depleted after spaceflight as well as *A. muris* (*BaiCD*), *E. massiliensis* (*BaiA*) and *B. pseudococcoides* (*BaiCD*), species which were significantly enriched after spaceflight (Supplementary file 2 and 3). *E. aldenensis* and *S. arabinosiphila* both contained the important *BaiCD* gene and were significantly enriched in mice after 29 days of spaceflight but significantly depleted after 56 days. Alongside significant enrichment of the rare 7α-dehydroxylating *E. muris*, these shifts suggest dynamic changes in secondary bile acid production and potential influence on the composition of the host bile acid pool^56,79^.

Differential abundance analysis inferred 52,370 genes were significantly different (FDR < 0.1) in abundance after 29 days and 37,068 genes after 56 days of spaceflight, which could be assigned to 2,811 and 2,572 unique KEGG ontology terms, respectively (Fig 4A). Over-representation analysis identified significant (FDR < 0.1) increases in pathways of interest related to fatty acid metabolism, bile acid metabolism, antimicrobial resistance and potential host interactions (ECM, carbohydrates and pathogenicity), after both 29 days and 56 days of spaceflight compared to ground controls (Fig 4BC). Taken together, these significant changes in metagenomic gene inventories, and specific bacterial species with well-characterised functions, due to spaceflight suggest gut microbiome changes which should influence lipid and bile acid homeostasis, and the immune system of the murine host.

**Figure 4.**
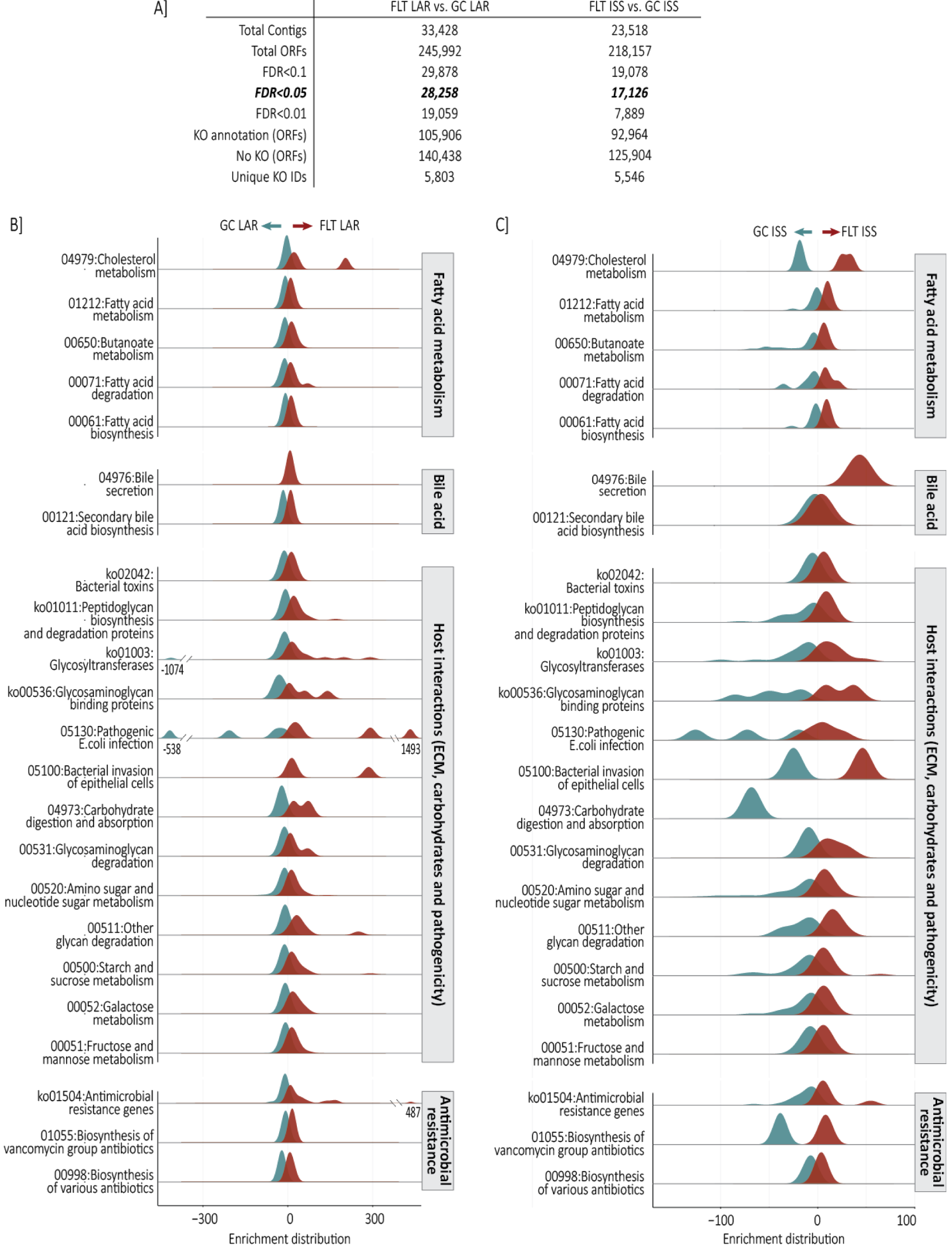
Metagenomic functional prediction. Summary statistics from metagenomics functional prediction (further detail in Supplementary file 4). B) GC_LAR vs FLT_LAR Over-representation analysis (ORA) of KEGG ontology (Brite, pathway and module) and C) GC_ISS vs FLT_ISS ORA KEGG ontology. Changes in fatty acid pathways included Fatty acid biosynthesis (00061), Fatty acid metabolism (01212), Fatty acid degradation (00071) and Butanoate metabolism (00650), including butyryl CoA:acetate CoA transferase (EC 2.8.3.8) and butyrate kinase (EC:2.7.2.7). Over-representation of bile acid metabolism was reflected in Bile secretion (04976) and Cholesterol metabolism (04979) and Secondary bile acid biosynthesis pathways, including bile salt hydrolase (*cbh*, EC:3.5.1.24) and 3-oxocholoyl-CoA 4-desaturase (*baiCD*, EC:1.3.1.115). Over-representation of the antimicrobial resistance was represented in Brite ontology Antimicrobial resistance genes (ko01504) and the pathways for beta-Lactam resistance (01501), Biosynthesis of various antibiotics (00998) and Biosynthesis of vancomycin group antibiotics (01055). The Brite ontology Bacterial toxins (ko02042) was over-represented, including tight junction interacting zona occludens toxin (K10954), as well as the pathways Pathogenic Escherichia coli infection (05130) and Bacterial invasion of epithelial cells (05100). Diverse carbohydrate metabolism and ECM interacting pathways were represented by Galactose metabolism (00052), Mannose type O-glycan biosynthesis (00515), Glycosaminoglycan degradation (00531), Other glycan degradation (00511), ECM-receptor interaction (04512) as well as the Brite ontology of Glycosaminoglycan binding proteins (ko00536), Peptidoglycan biosynthesis and degradation proteins (ko01011) and Glycosyltransferases (ko01003). These included putative Mucin-associated glycosyl hydrolases (GHs)^186^, GH2s: β-galactosidase (EC:3.2.1.23), β-mannosidase (EC:3.2.1.25), β-glucuronidase (EC:3.2.1.31), α-l-arabinofuranosidase (EC:3.2.1.55), β-xylosidase (EC:3.2.1.37), β-glucosidase (EC:3.2.1.21), GH20: β-hexosaminidase (EC:3.2.1.52), GH29: α-l-fucosidase (EC:3.2.1.51), and GH84: N-acetyl β-glucosaminidase (EC:3.2.1.52).

### 2.3 Spaceflight alters host intestinal gene expression

Faecal or serum fatty acid or bile acid concentrations were not measured within the Rodent Research 6 mission; however, host colon and liver gene expression were assessed from all four groups of mice, allowing host responses to spaceflight at the host-gut microbiome interface to be investigated.

Host intestinal gene expression revealed extensive significant (FDR <0.1) changes after 29 days and 56 days of spaceflight when compared to ground controls, including 4,613 differentially expressed (DE) genes between GC_LAR and FLT_LAR, and 4,476 DE genes between GC_ISS and FLT_ISS (Fig 5A-H; Supplementary document 1 and file 5). Of these, 43% and 44% were increased due to flight in LAR and ISS mice, respectively. Gene set enrichment analysis (GSEA) revealed consistent responses at a pathway level after 29 days and 56 days of spaceflight (Fig 5IJ), including immune suppression, dysregulation off cholesterol and bile acid, and extracellular matrix (ECM) remodelling.

**Figure 5.**
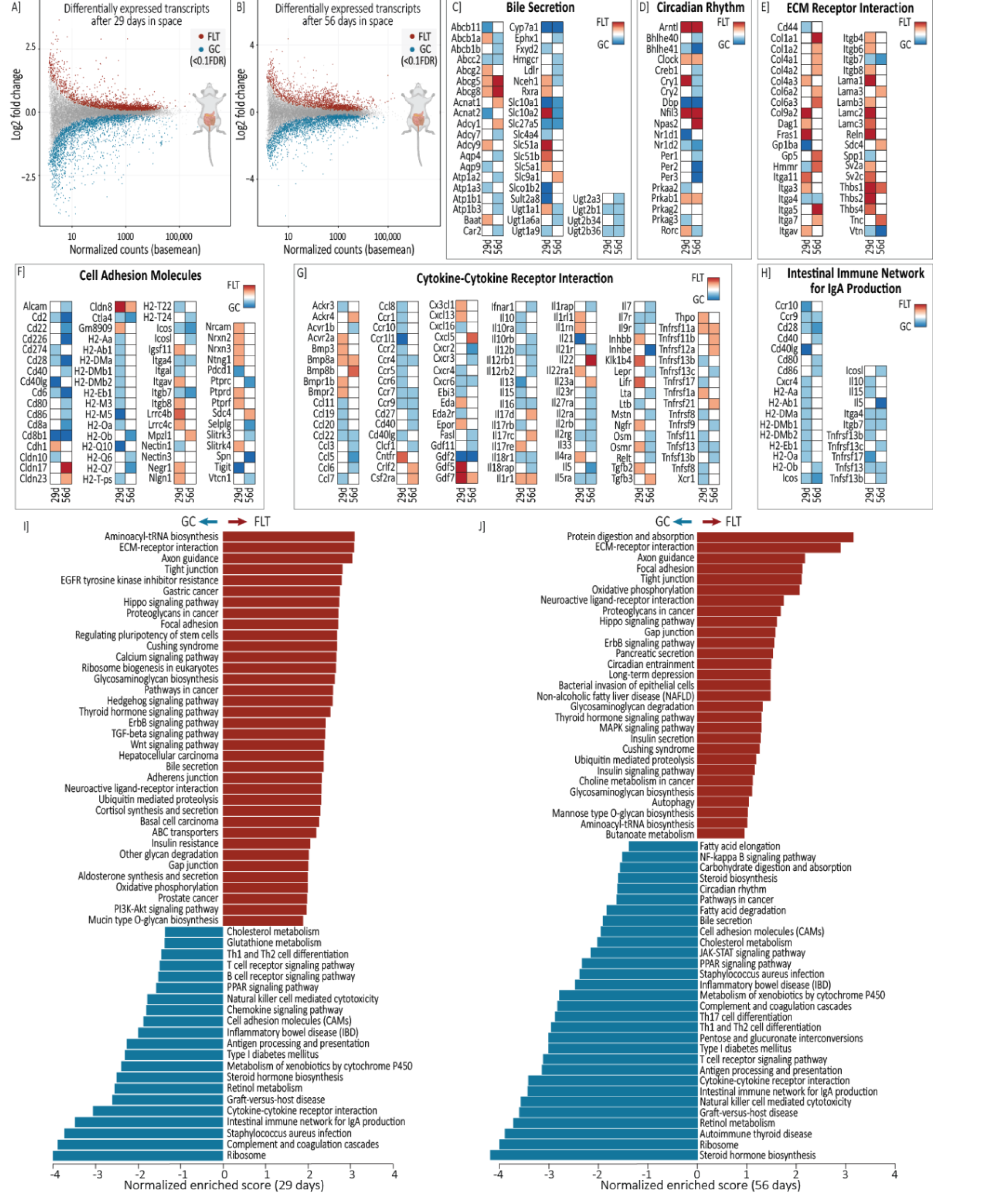
Micorbiome-host interface: spacelfight alters colon gene expression. MA plots showing A) GC_LAR vs FLT_LAR (29 days of spaceflight) and B) GC_ISS vs FLT_ISS (56 days of spaceflight) differentially expressed genes in the colon (FDR < 0.1). Differentially expressed gene from select KEGG pathways of interest, I) Significant Gene Set Enrichment Analysis (GSEA) 29 days (modified from WebGestalt^187^) and J) Significant GSEA 56 days spaceflight. Full DE gene list is available in Supplementary File 5 and select KEGG pathways of interest with differentially expressed gene highlighted are available in Supplementary document 1. Gene set enrichment analysis (GSEA) revealed consistent responses at a pathway level between 29 days and 56 days of spaceflight. This included widespread downregulation of the components of the intestinal immune system after spaceflight, including intestinal immune network for IgA production, antigen processing and presentation, Th1 and Th2 cell differentiation, PARR signalling metabolism of xenobiotics, *Staphylococcus aureus* infection, T cell receptor signalling, natural killer cell mediated cytotoxicity, graft-vs-host disease and cytokine-cytokine receptor interactions pathways, as well as downregulation of cholesterol pathways, including cholesterol metabolism and steroid hormone biosynthesis. Spaceflight also led to common upregulation of pathways associated to intestinal extracellular matrix (ECM) remodelling, including ECM-receptor interactions, focal adhesion, tight junction, gap junction pathways, and cortisol production represented through the Cushing syndrome pathway. The bile secretion pathway was significantly upregulated after 29 days of spaceflight, but downregulated after 56 days, suggesting bile acid dynamics should be explored at the gene level. Similarly, mucin type O-glycan biosynthesis, pathways in cancer and insulin resistance were only upregulated at the pathway level after 29 days of spaceflight, while bacterial invasion of epithelial cells, NAFLD, butonoate metabolism, insulin secretion and insulin signalling pathways were upregulated and the circadian rhythm pathway was downregulated after only 56 days of spaceflight.

#### 2.3.1 Intestinal bile acid and circadian rhythm gene expression

Significant microbiome alteration in some of the few well-characterised 7α-dehydroxylating bacterial species, including increases in *E. muris* after 29 days and 56 days of spaceflight (below detection in ground controls) as well as depletion of *C. scindens*, suggests secondary bile acid production and the bile acid pool is likely altered in the murine gut during spaceflight. In the host intestine, bile acids are passively absorbed or actively taken up through the apical membrane by the apical sodium-dependent bile acid transporter, ASBT (*Slc10A2*), bound to the cytosolic ileal bile acid binding protein, IBABP (*Fabp6*), and then transported across the basolateral membrane by organic solute transporters, Ostα and Ostβ (*Slc51A* and *Slc51B*), or glucuronidated by UGTs (such as *Ugt1a1)* and exported back to the lumen by multidrug resistance-associated protein 2, MRP2 (*Abcc2* transporter)^80,81^. Increases in intestinal bile acid also activate the farnesoid X receptor (FXR) and retinoid X receptor α (RXRα) heterodimer which regulates production and secretion of fibroblast growth factor *FGF15/19*, the negative feedback hormone which travels through portal circulation to bind hepatic FGFR4 receptors which suppress liver bile acid biosynthesis via inhibition of *Cyp7A1*^82^.

After 29 days of spaceflight, *Abcc2* was significantly downregulated, suggesting reduced BA export to the colon, while *Asbt*, *Ibabp, Ostα*, *Ostβ* and *Ugt1a1* were all significantly upregulated as well as *Rxrα* (not *Fxr*), *Fgf15/19* and *Fgfr4* (Fig 5C; Supplementary document 1 and file 5). Taken together, these changes suggest spaceflight led to an alteration in bile acid metabolism in the intestine which would lead to bile acid suppression of hepatic *Cyp7A1* and accumulation of cholesterol or hypercholesterolemia^83^. Interestingly, intestinal *Cyp7A1* expression was identified as significantly repressed after 29 days of spaceflight. Previous proteomic research in Biom-1M mice suggested a decrease in bile secretion during spaceflight^84^ while hepatic metabolite assessment of mice after spaceflight in the Space Shuttle Atlantis measured increased accumulation of cholate and taurodeoxycholate^16^. As the major cholesterol degradation mechanism in humans and mice is conversion to bile acids, cholesterol accumulation, alongside bile acid dysregulation and suppression of *Cyp71A,* should increase direct intestinal cholesterol excretion^85^. Supporting this extrapolation, both cholesterol excretion transporter genes, *Abcg5* and *Abcg8*, were significantly upregulated in the intestine after 29 days of spaceflight.

After 56 days, intestinal *Cyp7A1* was still significantly repressed and both *Abcg5* and *Abcg8* upregulated, but GSEA indicated a further shift in intestinal bile acid metabolism (Fig 5IJ; Supplementary document 1). This was underlined by significant downregulation of the apical bile acid transporter *Asbt* compared to ground controls as well as *Ntcp* (*Slc10a1*)^86^, indicating a switch to active reduction in bile acid uptake. Coinciding with this was significant increases in expression of *Lxrβ*, the liver x receptor gene expressed widely in different tissues, which can help prevent bile acid toxicity through induction of Abcg5 and Abcg8 mediated cholesterol excretion^87–90^.

In gene expression analysis of multiple tissues in mice after spaceflight, da Silveira et al.^18^ found that enrichment within the circadian rhythm pathway of the kidney, liver, eye, adrenal gland and various muscle tissues. Within the intestine here, the major clock genes, the circadian locomotor output cycles kaput (*Clock*) and brain and muscle ARNT-like protein-1 (*Bmal1*; *Arntl*) transcription factors, were significantly upregulated after 29 days and 56 days of spaceflight, and the neuronal PAS Domain Protein 2 (*Npas2*) was upregulated after 56 days (Fig 5D; Supplementary document 1 and file 5). These regulate the major clock-controlled genes reverse-erythroblastosis (Rev-Erbα and β) and retinoic acid receptor-related orphan receptors, including gamma (*Rorc*)^91^, as well as *Period* (*Per1, Per2 and Per3*), *Cryptochrome* (*Cry1* and *Cry2*) and basic Helix-Loop-Helix (bHLH) protein (*Dec1* and *Dec2*) genes^92,93^, all of which were significantly downregulated after 56 days of spaceflight except for *Rev-Erbα* (*Nr1d1*) (only downregulated after 29 days). These genes can feedback to inhibit *Clock/Npas2* and *Bmal1* as part of a feedback loop^94,95^, and also regulate other the clock controlled genes such as *DBP, HLF and TEF,* significantly downregulated, and *Nfil3 (E4BP4)*, significantly upregulated after 56 days of spaceflight. These spaceflight-associated changes in core clock genes, such as upregulation of *Bmal1*, *Arntl* and *Npas2,* largely agree with those found in murine muscle tissue after longer-term spaceflight which were characterised as similar to age-related gene expression on earth^96^.

These clock genes regulate nutrient absorption, gut motility, intestinal barrier function and immunity^92^ and have also been shown to require and interact with the microbiome, including in response to microbially-derived molecules, such as butyrate and bile acids, and in direct response to microbial associated molecular patterns (MAMPs)^53,92,97–99^. In addition to the widespread intestinal gene expression responses to bile acids, the major butyrate receptor free fatty acid receptor 2 (*Ffar2*; *Gpr43*)^100^ was significantly upregulated after 56 days of spaceflight. Similarly, the archetypal intestinal MAMP recognising mannose-binding lectin genes, *Mbl-1* and *Mbl-2*^101^, were significantly downregulated. Furthermore, Wang et al ^102^ reported host immune responses to microbial flagellin and lipopolysaccharide in the intestine increased expression of Il-23, and Il-22, leading to a downregulation of *Rev-Erb* and subsequent upregulation of *Nfil3*, which in turn can regulate clock-associated nutrient absorption and immunity. Here, mice followed this specific expression pattern after spaceflight (Supplementary file 5), implying the spaceflight associated microbiome alterations could have been recognised and influenced these changes in clock gene expression.

#### 2.3.2 Intestinal extracellular matrix remodelling and immune compromise during spaceflight

Changes in the extracellular matrix-receptor interactions pathway were underpinned by significant upregulation of collagens (*Col1a1*-*2*, *Col3a1*, *Col4a1*-*2*, *Col5a1*-*2*, *Col5a3*, *Col6a2*-*3*, *Col12a1*, *Col26a1*), laminins (*Lama3)*, thrombospondin (*Thbs1*) and tenascins (*Tnc*) in mice after 29 days of spaceflight, which became more pronounced after 56 days of spaceflight (Fig 5E; Supplementary file 5). This coincided with upregulation of integrin (*Itga5* and *Itga7*) and matrix glycoprotein (*Sdc4* and *Gp5*) receptors, collectively implying extensive extracellular matrix remodelling during spaceflight. The ECM and mucosal collagen scaffold in particular are known to be shaped by microbiota^103^, as are mucins, which make up the intestinal mucus layer and have a dynamic relationship with commensal bacteria as well as serving as a critical barrier against colonisation by pathogenic bacteria^104^. Significant increase in the secretory mucin gene *Muc2* and significant decreases in the membrane bound *Muc3* and mucosal pentraxin 1 (Mptx1), three of the mostly highly abundant transcripts in the murine intestine here, were observed after 56 days of spaceflight, suggesting alterations to mucin within the intestinal lumen in direct contact with microbiota. Mucin 2 (*Muc2*) is well characterised as regulated by intestinal bacteria, with O-glycans serving as nutrients and adhesion sites for microbiota^105^, but are also differentially expressed in response to pathogens^104^, including *Trichinella*^106^, identified here.

Extensive changes to mucosa were indicated by widespread downregulation of cell adhesion molecules during spaceflight, including downregulation of *CD8a* and *CD8b1* genes, genes encoding costimulatory molecules *CD2*,*6*,*80*,*86*,*40* and *ICOSL* within intestinal antigen presenting cells as well as their T cell activating binding partners *CD48*,*166*,*28* and *ICOS*^107–109^ (Fig 5G; Supplementary file 5). Cytokine genes, such as the chemokine Ccl22 and receptor Ccr4 involved in the intestinal immune response to enteric bacterial pathogens in murine mucosa^110^, were also uniformly downregulated (Fig 5F), alongside others^111–113^: *Ccl3*,*5*,*6* and *22, Ccr4*,*7 and 9, Cxcr2,3 and 6*, *Il-5*,*7*,*12* and *16*, and *Il-2r*,*5r*,*7r*,*10r*,*12r*,*18r*,*21r*,*23r* and *27r.* An exception to this pattern of cytokine downregulation was upregulation IL-23 and IL-22, which interact with circadian regulation^95^, and specific members of the mucosal homeostasis critical interleukin 17 family^114^, *Il-17d,* which promotes pathogenicity during infection through suppression of CD8+ T cells^115^, and *Il-17rc*, which increases expression during compromised epithelial barrier integrity (wounding)^116^.

These expression profiles, alongside consistent downregulation of genes within the Intestinal IgA pathway (fig 5H), suggest suppression of immunity and widespread tissue remodelling at the host-gut microbiome interface in mice after spaceflight. This agrees with reports of reduced cytokine production in mice after simulated microgravity^117^, immune dysfunction in splenic tissue of mice after 13 days of spaceflight on the Space Shuttle Atlantis^118^ and in astronauts, alongside increases in plasma cortisol concentration which reached Cushing syndrome levels, during spaceflight^119^. Taken together, this provides an insight into the role the host-gut-microbiome interface might play in the current broad consensus of immune dysregulation in spaceflight environments^120^.

### 2.4 Spaceflight alters gene expression in the liver

Hepatic gene differential expression analysis comparing mice after 29 days and 56 days in space to their relative ground controls identified 4,029 DE genes and 4,068 DE genes, respectively (FDR < 0.1; Fig 6AH; Supplementary document 1 and file 5). Of these, 48 % and 49% were increased due to 29 days and 56 days of spaceflight, respectively. GSEA of liver tissue responses also revealed highly consistent responses at the pathway level to 29 and 56 days of spaceflight (Fig 6IJ), including disruption of bile acid and energy metabolism.

**Figure 6.**
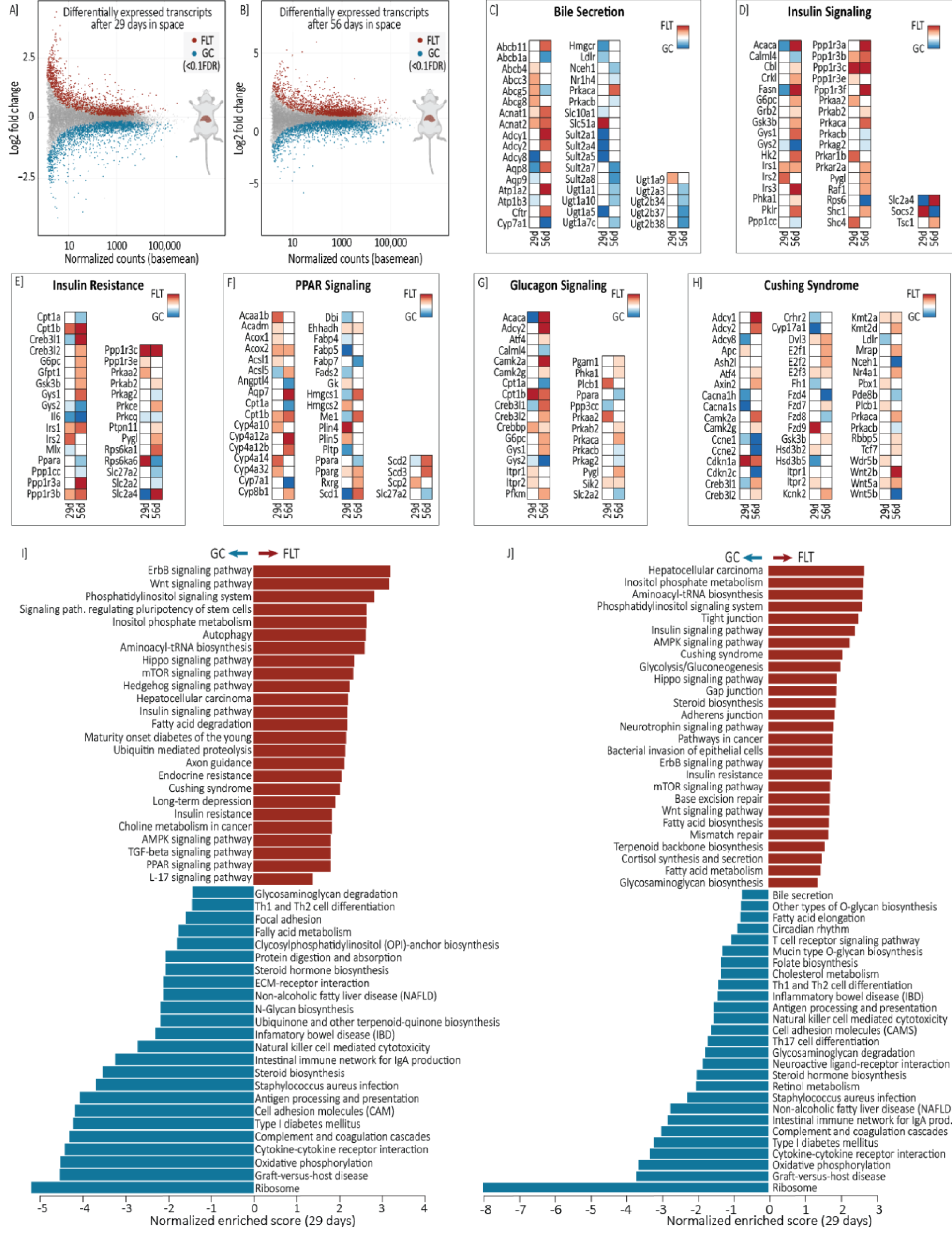
Microbiome-host metabolism: spaceflight alters liver gene expression. MA plots showing A) GC_LAR vs FLT_LAR (29 days of spaceflight) and B) GC_ISS vs FLT_ISS (56 days of spaceflight) differentially expressed genes in the liver (FDR < 0.1). Differentially expressed gene from select KEGG pathways of interest, I) Significant Gene Set Enrichment Analysis (GSEA) 29 days (modified from WebGestalt^187^), and J) Significant GSEA 56 days spaceflight. Full DE gene list is available in Supplementary File 5 and select KEGG pathways of interest with differentially expressed gene highlighted are available in Supplementary document 1. GSEA of liver tissue responses also revealed highly consistent responses at the pathway level to 29 and 56 days of spaceflight. These comprised downregulation of immune response pathways, similar to those seen in the intestine, as well as steroid metabolism, type I diabetes mellitus, inflammatory bowel disease and NAFLD. Spaceflight also led to common upregulation of insulin resistance, Hippo signalling, inositol phosphate metabolism, Cushing syndrome and hepatocellular cancer pathways at both 29 and 56 days. Certain pathways were different over time. After 29 days of spaceflight, long-term depression and maturity onset diabetes of the young pathways were upregulated, whereas after 56 days, bile secretion and circadian rhythm were downregulated, while glycolysis/gluconeogenesis pathway were upregulated.

#### 2.4.1 Bile acid disruption underlies hepatic cholesterol accumulation

The most abundant intestinal transcripts in mice after 56 days of spaceflight, differentially expressed and representing 8% and 9% of normalised counts, were from the non-coding RNA *Rn7s1* and *Rn7s2* genes (7S RNA 1 and 2; Supplementary file 5), respectively, which were recently characterised as inhibitors of global mitochondrial transcription in mammals^121^. This is consistent with the mitochondrial dysfunction highlighted as characteristic of spaceflight pathogenicity in recent multiomic analysis performed by de Silveira et at^18^, who also characterised compromised liver function in mice and astronauts after spaceflight compared ground controls, including upregulation of *Fgf21*, a negative repressor of bile synthesis^122^, and accumulation of total cholesterol (higher low-density lipoprotein cholesterol but decreased high-density lipoprotein cholesterol). Here, another potent repressor of *Cyp7A1* bile synthesis, fibroblast growth factor 1 (*fgf1*)^123^, was upregulated in the liver of mice after 29 days of spaceflight, and the more well known *Fgf21*^16^ was significantly upregulated in the liver after both 29 days and 56 days of spaceflight.

Within the liver, Sterol 14-demethylase (Cyp51) catalyses the transformation of lanosterol into cholesterol and Cyp7A1 is then the first (and rate-limiting) enzymatic step in transformation of cholesterol in primary bile acid biosynthesis, which itself is strictly inhibited by bile acid concentrations^124^. The production of bile salts is then a stepwise transformation process dependent on gene expression of 3 beta-hydroxysteroid dehydrogenase type 7 (*Hsd3b7*) gene, cholic acid-specific *Cyp8b1* gene and acyl-Coenzyme A oxidase 2 (*Acox2*) before conjugation by *Baat*, *Acnat1* and *Acnat2*, and subsequent excretion in the bile duct by bile salt exporter pump (BSEP)^125,126^. *Cyp51*, *Cyp8b1* and *Acox2* genes were significantly upregulated and *Cyp7a1* and *Fxr* (*Nr1h4*) were significantly downregulated in the liver of mice after 56 days on the ISS (Fig 6C; Supplementary document 1 and file 5). The cholesterol transporter genes *Abcg5* and *Abcg8* (intestinally upregulated after 29 and 56 days of spaceflight) were upregulated in the liver after 29 days, but no longer significantly upregulated after 56 days, and *Abcg8* was significantly downregulated. This reduction is surprising given the uniform accumulation of cholesterol observed after extended spaceflight; in contrast, *Bsep* and the bile acid conjugating acyl-coenzyme A:amino acid N-acyltransferase 1 and 2 (*Acnat1* and *Acnat2; Baatp1/2*) genes were significantly upregulated. Taken together, these expression profiles suggest hepatic accumulation of cholesterol, characteristic of glucose and lipid metabolic dysruption^97^, and describe a subsequent increase in the production bile acids in the liver, preferentially cholic acid, their subsequent conjugation and export as bile salts, with the seemingly contradictory reduction in C*yp7A1* consistent with feedback inhibition.

The other major mechanism for detoxification during cholesterol and bile acid accumulation in the liver is sulfonation of bile acids, the transfer of a sulfonate group to a hydroxyl (OH) by a subfamily of cytosolic sulfotransferases (*Sult2a* genes) which increases their solubility, decreases enterohepatic recirculation, and increase excretion^127^. After 29 days of spaceflight, hepatic *Sult2a1*, *Sult2a4* and *Sult2a5* were downregulated, which shifted to downregulation of *Sult2a7* and *Sult2a8* after 56days (Supplementary file 5). In humans, bile acid sulfonation is catalysed by Sult2a1, which sulfonates the 3-OH of bile acids. In contrast, mice have 8 *Sult2a* genes, with *Sult2a1-6* sharing close homology to *Sult2a1* but *Sult2a8* being recently characterised as having major function in sulfonating 7α-OH of bile acids^128,129^, of particular relevance here due to spaceflight microbiome changes in 7α-dehydroxylating *E. muris* and *C. scindens*. Notably, during acute phase immune responses, alterations to fatty acid, cholesterol, and bile acid metabolism, *Sult2a1* is known to be suppressed by cholesterol and bile acid regulating nuclear *Fxr* (*Nr1h4*) and *Car* (*Nr1i3*) nuclear receptors^130^, the latter of which also regulates bile acid responsive transporter gene *Mdr1* (ABCB1)^131^, all three of which were downregulated after 56days of spaceflight (Fig 6c) and provide further evidence of bile acid dysregulation and toxic stress consistent with extensive disruption of the gut-liver axis^132^.

*2.4.2 Energy homeostasis disruption after spaceflight*

Beheshti et al.^17^ observed significant depletion in Cyp7A1 protein levels in mice after spaceflight (RR1 and RR3), alongside disruption in glucose and lipid metabolism^133^ as well as NAFLD ^134^. Here, pathways related to energy homeostasis consistently altered in gene expression due to spaceflight after 29 and 56 days included enrichment of fatty acid degradation, insulin signalling and insulin resistance (Fig 6DE; Supplementary document 1 and File 5). The highest relative abundance (CPM) of transcripts significantly increased in the liver of mice after 56 days of spaceflight included fatty acid synthase (*Fasn*), the liver fat accumulation-associated carbonic anhydrase 3 (*Car3*)^135^, and the rate limiting enzyme for fatty acid desaturation, Stearoyl-CoA desaturase (SCD), recently identified as a key role at the crossroads of immune response and lipid metabolism through interplay with PPARγ^136^, also significantly upregulated here (fig 6F). Glucose metabolism was also disrupted by spaceflight as the glucose transporter *Glut2* (*Slc2a2*) gene, required for glucose-stimulated insulin secretion^137^, and glycogen synthase 2 (*Gys2*) gene, the rate limiting enzyme for glycogenesis^138^, were downregulated in the liver (and intestine) after both 29 days and 56 days of spaceflight. The free fatty acid and glycolysis regulating *PPARα* was also downregulated, and liver glycogen phosphorylase (*Pygl*) and glycogen synthase kinase 3 beta (*Gsk3β*) were significantly upregulated after 56 days of spaceflight (Fig 6G). Taken together, the indicated decrease in glycogen synthesis and increase in glycogenolysis is characteristic of insulin resistance leading to the elevated fasting plasma glucose to pre-diabetic levels previously observed in crew of the Mars500 analogue mission and during spaceflight^139–141^. Interestingly, in light of spaceflight induced changes to gut microbiota, upregulation of *Gsk3β* is also known to be activated by microbial-associated molecular patterns^142^ and promotes acute liver failure through inhibition of the PPARα pathway^143^.

Other pathways enriched in the liver after spaceflight included Cushing Syndrome, hypercortisolism consistent with elevation of cholesterol levels, and hepatocellular cancer pathways. Notably, claudins, which were largely downregulated in the intestine where they are commonly expressed within tight junctions, were upregulated in liver tissue after 56 days of spaceflight (Supplementary File 5), including highly significant and high relative abundance increases in *CLDN1* and *CLDN2*. So-called non-tight junction claudins have only recently been experimentally explored and, in hepatic cells, CLDN1 is implicated in hepatocellular carcinoma (HCC)^144^. More broadly, significant upregulation of *Tgf-α* and genes involved in complex Tgf-β signalling (*Tgfb2*, *Tgfb3*, *Tab2*, *Tgfbrap*, *Smad3*), alongside other markers (*Dapk2*, *Vegfa*, *Dvl3*)^145,146^ are associated to HCC, as well as immune suppression through the cyclin-dependent kinase inhibitor 1A (*p21^cip^*^1^; *Cdkn1a*)^147^, also significantly upregulated after 29 and 56 days of spaceflight. Previous gene expression analysis of mice exposed to high-energy ion particle radiation to simulate exposure to Galactic Cosmic Rays reported induction of spontaneous HCC^146^. These prominent transcriptomic shifts after 56 days of spaceflight, if reflected in longer term studies in humans, could represent a serious health concern.

### 2.5 Conclusions

Through metagenomic assessment of the murine gut microbiome, significant spaceflight-associated changes in bacteria linked to bile and fatty acid metabolism were identified. These changes in relative abundance were largely consistent in two groups of mice after spaceflight when compared to different on-Earth control groups at different timepoints as well as when using distinct metagenomic methodologies.

The microbiome changes coincided with substantial changes to gene expression at the host-gut microbiome interface which are critical to barrier function, microbe interactions and bile acid transport in the intestine. These interactions suggest disruption of the signals, metabolites, and immune factors exchanged across the gut-liver axis which are likely to drive glucose and lipid dysregulation. Collectively, these multiomic findings suggest host-gut microbiome interactions during spaceflight are likely to underly widespread changes to host physiology which could pose a risk to health.

## 3 Funding

NB would like to acknowledge support from the University College Dublin Ad Astra program. Genelab is funded by the Space Biology Program (Science Mission Directorate, Biological and Physical Sciences Division) of the National Aeronautics and Space Administration. Open access was funded by University College Dublin.

## 4. Acknowledgements

This research was generated by the NASA GeneLab Analysis Working Group for Microbes. The team would like to acknowledge the GeneLab for School (GL4S) team for supporting young scientists.

## 5 Supplementary Files

Supplementary document 1 – includes study limitations and future perspectives, and additional supplementary figures detailing mouse carcass weights, extended amplicon processing statistics, metagenomics processing statistics and host differential expression pathway maps.

Supplementary file 1 – RR6 amplicon-based metagenomics Supplementary file 2 – RR6 WMS processing summary Supplementary file 3 – RR6 WMS-based taxonomy Supplementary file 4 – RR6 WMS-based function Supplementary file 5 – RR6 host transcriptomics

## 6 Methods

### 6.1 Experimental design

Thirty-two 32-weeks-old female C57BL/6NTac mice were split into four treatment groups: flight ISS (FLT_ISS, n=7), ground control for ISS (GC_ISS, n=9), flight live animal return (FLT_LAR, n=9) and ground control for live animal return (GC_LAR, n=7) (Fig 1A). FLT_ISS and FLT_LAR mice were launched on SpaceX-13 and transferred to the rodent research habitat on the ISS whereas their matched ground controls, GC_ISS and GC_LAR, were kept in identical rodent habitats at the Kennedy Space Centre. Diet (LabDiet Rodent 5001) and deionized autoclaved water were provided *ad libitum*, and a 12:12 hr dark/light cycle maintained. After 29 days of flight onboard the ISS, FLT_LAR mice were returned to earth as part of the Live Animal Return and sacrificed alongside GC_LAR using common processing at ages of 41 weeks old. FLT_ISS mice were sacrificed after 53-56 days of flight onboard the ISS at the same time as GC_ISS mice at the Kennedy Space Centre at 44 weeks old using a common timeline and methodology.

During this period in the Destiny module (US laboratory) on the ISS, the mice were exposed to an average daily 165.8 µGy d^-1^ Galactic Cosmic Ray (GCR) dose and 117.3 µGy d^-1^ South Atlantic Anomaly (energetic protons) dose (data provided by Ames Life Sciences Data Archive - ALSDA). This is in line with standard range of exposure on the ISS^148^, and represents around a 100% increase to common exposure on earth. The temperature, relative humidity and elevated carbon dioxide levels on the ISS were mimicked in the ground control rodent habitats at the Kennedy Space over the 56 days of spaceflight, so were not significantly different (t-test, p>0.05) between flight and ground controls, and averaged 22.75 (±0.35) °C, 41.49 (±2.28) % and 3,219 (±340) CO_2_ ppm, respectively.

### 6.2 Murine colon and liver RNA, and intestinal metagenomic DNA sampling and sequencing

#### 6.2.1 DNA extraction

DNA was extracted using the Maxwell RSC Purefood GMO and Authentication Kit (Promega, Madison, WI) (<u>OSD-249</u>). Half of a frozen faecal pellet was placed into a tube with 940 uL CTAB solution and homogenized using tissue homogenizing bead mix (Navy RINO Lysis, Next Advance) on Bullet Blender Gold 24 (Next Advance) for 4 minutes at 4°C. Homogenates were centrifuged for 3 minutes at 10°C and 21,000 g to deflate foam. The supernatant from each sample was then used to isolate and purify DNA following the manufacturer’s protocol. DNA was eluted in 105 µL RNAse free H_2_O and was further cleaned using OneStep PCR Inhibitor Removal Kit (Zymo Research). Concentrations for all DNA samples were measured using Qubit 3.0 Fluorometer (Thermo Fisher Scientific, Waltham, MA) with a Qubit DNA HS kit. DNA quality and size were assessed using an Agilent 4200 TapeStation with a gDNA ScreenTape Kit (Agilent Technologies, Santa Clara, CA).

#### 6.2.2 16S rRNA gene amplification

DNA library preparation was performed by the Genome Research Core (GRC) at the University of Illinois at Chicago. 10 ng of genomic DNA was used as input to a two-stage PCR amplification protocol^149,150^. In the first stage, primers 515F/806R (Earth Microbiome Project) containing Fluidigm ‘Common Sequence’ linkers (CS1 and CS2) were used to amplify gDNA. In the second stage, Fluidigm AccessArray barcoded primers were used to amplify PCR products from the first stage and incorporate Illumina sequencing adapters and a sample barcode. Sequencing was performed on an Illumina MiniSeq mid-output flow cell, employing paired-end 2×153 base reads.

#### 6.2.3 Whole metagenome sequencing

Whole metagenome sequence libraries were prepared using an Illumina Nextera DNA Flex Library Prep kit (Illumina, San Diego, CA) according to the manufacturer’s instructions. Input DNA was approximately 100 ng per reaction, and five cycles of PCR were performed. Index adapters used were IDT for Illumina, 96-well Nextera Flex Dual Index Adapters, set A. Library fragment sizes (approximately 550 bp) were assessed using an Agilent 4200 TapeStation with D1000 DNA ScreenTapes (Agilent Technologies, Santa Clara, CA). Pooled library concentration was measured with a KAPA Library quantification kit (Roche, Wilmington, MA). Library quality control was performed on an Illumina iSeq100 sequencer (Illumina, San Diego, CA). Whole metagenome shotgun sequencing was performed on an Illumina NovaSeq6000 instrument with a 500-cycle SP flow cell. Library preparation and sequencing were performed by the GeneLab Sample Processing Lab (NASA Ames Research Center).

#### 6.2.4 RNA extraction and sequencing

RNA was extracted from mouse tissue samples using an AllPrep DNA/RNA Mini Kit (Qiagen, Valencia, CA). Homogenization buffer for RNA purification was made by adding 1:100 beta-mercaptoethanol to Buffer RLT (Qiagen, Valencia, CA) and kept on ice until use. Approximately 30 mg of frozen colon (<u>OSD-247</u>) or liver (<u>OSD-245</u>) tissue was isolated using a scalpel, weighed and immediately placed in 600 uL of the Buffer RLT solution. Homogenization was performed using tissue homogenizing bead mix (Zirconium Oxide 2.0mm Beads, Next Advance) on Bullet Blender Gold 24 (Next Advance) for 5 minutes at 4°C. Homogenates were centrifuged for 3 minutes at RT and 14,000 g to remove cell debris. The supernatant from each sample was then used to isolate and purify RNA following the manufacturer’s protocol. RNA was eluted in 50 µL RNAse free H_2_O. Concentrations for all RNA samples were measured using a Qubit 3.0 Fluorometer (Thermo Fisher Scientific, Waltham, MA). RNA quality was assessed using an Agilent 2100 Bioanalyzer with an RNA 6000 Nano Kit or RNA 6000 Pico Kit (Agilent Technologies, Santa Clara, CA). ERCC ExFold RNA Spike-In Mixes (Thermo Fisher Scientific, Waltham, MA Cat 4456739, v92) at 1:100 dilution of either Mix 1 or Mix 2 were added on the day of library prep at the concentrations suggested by the manufacturer’s protocol.

Ribosomal RNA depletion was performed using an Illumina TruSeq Stranded Total RNA Library Prep Gold kit. Input RNA amounts were approximately 500 ng; RNA RIN values were >4. Index adapters were 1.5 µM (IDT, 384-well xGen Dual Index UMI Adapters). 15 cycles of PCR were performed. Library fragment sizes (approximately 300 bp) were assessed using an Agilent 4200 TapeStation with a D1000 DNA ScreenTape (Agilent Technologies, Santa Clara, CA). Pooled library concentration was measured by Universal qPCR Master Mix (Kapa Biosystems, Wilmington, MA). Library quality control was performed on an Illumina iSeq100 sequencer (Illumina, San Diego, CA). Whole metagenome sequencing was performed on an Illumina NovaSeq6000 instrument with a 500-cycle SP flow cell. Library preparation and sequencing were performed by the GeneLab Sample Processing Lab (NASA Ames Research Center).

### 6.3 Bioinformatics

#### 6.3.1 16S rRNA gene barcoding

Amplicon sequence reads were processed and annotated using Anchor^28,35,139,151,152^. Exact sequence variants (ESV) were identified in place of operational taxonomic units (OTUs)^153,154^. Sequences were aligned and dereplicated using Mothur^155^ and a count threshold parameter of 96. Annotation at family, genus or species-level used BLASTn criteria of >99% identity and coverage to the NCBI 16S curated and NCBI nr/nt databases (January 2022 versions). Differentially abundant ESVs were manually assessed for quality. When the highest identity/coverage was shared amongst multiple different references, all annotations were retained and reported.

Differential abundance analysis was performed using DESeq2^156,157^, which performs well with sparse data and uneven library sizes^158^. Sparsity and count thresholds were applied whereby an ESV count in a single sample was required to be <90% of the count in all samples, and ESV counts were required to be >0 in at least 3 samples from the same group^35^. A false discovery rate (FDR; Benjamini-Hochberg procedure) <0.1 correction was applied^159^.

#### 6.3.2 Whole metagenome sequencing co-assembly and annotation

Quality control used Trim Galore! (v0.6.6)^160^, a wrapper script to automate quality and adapter trimming as well as quality control. Trim Galore is based on cutadapt (v2.10)^161^ and fastqc (v0.11.5)^162^. Trim Galore! PARAMETERS: --trim-n --max_n 0 --paired --retain_unpaired --phred33 --length 75 -q 5 --stringency 1 -e 0.1 -j 1. BBMAP^163^ was used to remove potential contamination from human using the masked version of hg19 human assembly. To remove redundancy in read dataset and reduce the computational load, reads were normalized using ORNA^164^ with the following parameters: -sorting 1 -base 1.7 -kmer 21.

MEGAHIT v1.2.9^165^ was used to assemble reads from all samples into one co-assembly using *meta-large* option. Kallisto (v0.46.2)^166^ expectation maximization algorithm was used to complete metagenomics read assignment and infer contig abundance^167^. Prodigal (v2.6.3)^168^ was used with the option *meta* to predict open reading frames (ORFs) and BLAST v2.3.0 ^169^ was used to annotate contigs sequence.

To assign contig taxonomy, a first alignment iteration was run using full contig lengths against the NCBI nr/nt database (January 2022) and Reference Viral Database (RVDB v v25.0). To further resolve nucleotide taxonomic annotation, a second alignment was run against all databases which included selected genomes (additional 1148 sequences) from NCBI refseq informed by first iteration. BLASTn was run using the following parameters: -evalue 1e-50 -word_size 128 -perc_identity 97. Contig alignment scores were compared between the three databases and the best bitscore was selected as the best alignment for a given contig. Descriptive statistics were also provided for contigs with a common species annotation that had an average alignment identity >97%, total alignment length > 2000nt and an average query coverage >20%. To validate ESV sequences using the metagenomics *de novo* assembly, ESVs were aligned to WMS contigs using BLASTn.

To annotate genes, three protein databases (NCBI nr, UniProtKB Swiss-Prot, and TrEMBL; January 2022) were searched using the translated sequences of the predicted proteins. BLASTx was run with the following parameters: -evalue 1e-10 -word_size 6 -threshold 21. Alignment scores were compared between the three databases and the best bitscore was selected as the best alignment for a given orf. GO, pfam, PANTHER, EMBL, InterPro, HAMAP, TIGRFAMs, STRING, HOGENOM, SUPFAM terms were mined from UniProtKB database. Amino acid sequences were used as input in the GhostKOALA webserver^170^ to add functional genes and pathways information. KEGG functional and taxonomic annotation was retrieved using complete and incomplete pathways. *Extibacter muris* strain DSM28560 bile acid-inducible operon sequence (baiBCDEFGHI)^62^, from were manually added to default KEGG database. One bai sequence did not have a KEGG term associated to it (baiG MFS transporter; bile acid transporter) and a temporary KEGG term was assigned to it (K9999).

#### 6.3.3 Metagenome assembled genomes (MAGs)

Using metagenome co-assembly from 3.4.2, genome binning was performed using MetaBAT2^171^. Genome quality estimation of all bins was performed using CheckM (version v1.1.6)^172^. Taxonomic classification was performed with Bin Annotation Tool (BAT) a pipeline for the taxonomic classification of metagenome assembled genomes^173^.

#### 6.3.4 Murine transcriptome reference mapping

Mouse liver and colon RNA-Seq reads were processed and assembled following NASA GeneLab consensus pipeline, as described previously^174^.

## 7 Statistical Analysis

### 7.1 Alpha and beta diversity

To estimate and compare microbial richness within samples, alpha diversity was measured using diversity indices using Phyloseq R library^175^ and was compared between groups with t-tests (parametric) or Mann-Whitney U (non-parametric) tests. Unsupervised multivariate analysis (ordination) was performed using Principal Coordinate Analysis (PCoA) with normalized counts (Supplementary file) while constrained ordination was based on distance-based Canonical Correspondence Analysis (CCA). Significance of constraints were assessed using ANOVA-like permutation testing for CCA (anova.cca). Vegan R library^176^ was used to conduct these analyses, statistics, and to produce graphs and draw dispersion ellipses. As an exploratory visualization of annotated WMS contigs, Uniform Manifold Approximation and Projection (UMAP) was used to reduce the dimensionality of beta diversity WMS contig count matrices. CPM normalized counts of differentially abundant species (30 in each comparison) were selected as input and umap function from the umap R package (v 0.2.10) was used for each comparison with default parameters.

### 7.2 Differential abundance/expression analysis

Prior to differential abundance analysis, sparsity and count thresholds were applied whereby an ESV/contig/transcript count in a single sample must be <90% of the count across all samples and ESV/contig occurrence must be at least ≥3 in samples within the same design factor.

Differential abundance (or expression) analysis was performed using DESeq2^177^ based on pre-processed raw abundance of ESVs/contigs/ORFs/transcripts. A false discovery rate (FDR; Benjamini-Hochberg procedure) < 0.1 was applied for statistical significance^156^. Missingness is a known challenge for negative binomial regression models (such as used in DESeq2) when analyzing zero-inflated abundance tables^178,179^, contigs with an absolute zero across all replicated samples belonging to a same factor were assumed to be structural zeros and flagged as significantly differentially abundant. To address conservative p-value distribution^180^ of RNA-Seq differential expression analysis, local FDR values were computed from DESeq2 *p*-values using fdrtool (v1.2.17)^181^ R library.

### 7.3 Functional enrichment analysis

Ghostkoala output was organized into a gene count table using WMS ORF raw count table and used as input for over-representation analysis (ORA) of WMS data. ORA was used to statistically test the overlap between DA ORFs (FDR < 0.1) and a geneset using pathways of interest (Supplementary file 4). p-values were calculated using a hypergeometric test using clusterProfiler (v4.7.1.003) R library^182^.

Gene-set enrichment analysis (GSEA) of RNASeq data was performed on the Webgestalt^183^ platform using the entire gene list, rank-ordered combining significance and effect size from DESeq2 differential expression analysis, i.e. log2(FC)*-log(pValue)^184^. Gene symbols were inferred from assembly transcripts using org.Mm.eg.db (v3.16)^185^ R annotation library.

## Notes

https://osdr.nasa.gov/bio/repo/data/studies/OSD-249

### Competing Interest Statement

The authors have declared no competing interest.

