## Supplementary document for "Spaceflight alters host-gut microbiota interactions"

#### 1 Study limitations and future perspectives

A challenge for biological experimentation conducted in space is low replication, compared to model studies using mice on Earth. The use of distinct metagenomic methodologies, 16S rRNA gene barcoding and *de novo* whole metagenome sequencing, helped provide technical confidence in the spaceflight-associated changes in microbiome species detected here; however, it is likely that low n reduced statistical power and that important microbiome associations were missed, particularly those in low abundance bacteria. It is important that new systems being developed for commercial low earth orbit destinations (CLDs), Gateway and Lunar Base Camp as Artemis progresses are increasing capacity of experimental habitats and growth facilities for replicate numbers suitable for multiomic science in order to maximise future discoveries as well as enable future interventional microbiome studies.

Another weakness of this research is the lack of truly independent cohorts of mice, as Liver Animal Return (29 days spaceflight) and ISS (56 days spaceflight), and their respective ground controls, derived from the same initial cohort of mice. Common data gathering strategies would enable high resolution multiomic data to be directly compared across cohorts. Integrating metagenomic and host transcriptomic data here provided a highly complementary picture of the multiomic response to spaceflight and is a strength of an Open Science culture and public multiomic resources such as GeneLab. The addition of untargeted serum and faecal metabolomics would have further strengthened this research and future studies should aim to quantitatively profile the specific deconjugated, dehydroxylated, oxidated, and epimerized bile acids that are driving change in the system. Alongside metatranscriptomic analysis of the gut microbiome, this would represent an important next step likely to help clarify the directionality of the host-gut microbiome responses associated to spaceflight pathology.

#### 2 Identification of *Trichinella* in all murine faecal samples

The identification of parasitic worms (*T. nativa*) across all mice samples could indicate widespread trichinosis potentially linked to the built-environment of rodent laboratory settings[51-53]. The life cycle of *Trichinella* includes an enteral phase where larval worms within the small intestine penetrate epithelial cells, activating Th1 type immune responses and inflammation before maturing, mating, and producing larva. These larva can then induce a dominant Th2 type (helminth) immune responses in hosts aimed towards expulsing the parasite[54]. During the parental phase, larva circulate through the lymphatic and blood systems to enter and damage skeletal muscle. Within the context of spaceflight, exploration of changes in *Trichinella* relative abundance or gene expression are important due to potential interactions with mammalian muscle development, which is compromised in crew during to long-term confinement despite exercise intervention in ground-based experiments, such as MARS500[55,56], and as a consequence of microgravity in space.

#### 3 16S rRNA gene amplification and whole metagenome sequencing comparison of DA species

To validate the putative species annotation from amplicon analysis, ESVs annotated as species were aligned to WMS contigs. These ESVs could be aligned to WMS contigs at 99.3% ANI and 28/35 species identified using amplicon analysis were independently identified within the WMS co-assembly (including improved resolution of 10 ambiguous calls using WMS). Three of the seven species not identified in WMS were reclassified to a different species within the same genus based on WMS findings (database errors), two were present in WMS and likely represents a strain <97% identity to a known genome (one resolved during analyses as *Acinetobacter courvalinii*, genome published 2023), while three remaining species could represent limited WMS depth or regent contaminants unique to amplification (Supplementary file 3).

#### 4 Observing low abundance bacteria are important for statistical microbiome analysis

The majority of contigs were relatively short in length (Supplementary file 3) and associated to a small number of species with low relative abundance. Interestingly, contigs from some of these low abundance species were strictly associated to spaceflight, being present in mice after 29 and 56 days of spaceflight, and absent or below detection (structural zeros[51]) in their respective ground controls, including *D. welbionis*, *E. muris* and, in FLT_ISS samples, *G. tenuis* (Supplementary file 3). A limitation to this study was the moderate sequencing depth, with an average 8.7 M reads per sample. Minimum WMS depth requirements to capture most metagenomic gene content has been estimated as high as 80M reads in complex samples, and 200M reads may be insufficient to capture all genetic diversity[52]. Consequently, while more shallow WMS can be informative[53], the assembly and quantification across biological replicates of metagenome assembled genomes (MAGs) from low abundance species is challenging here due to insufficient sequencing coverage (Supplementary file 2). Recent high depth sequencing and culturing efforts are providing insight into the presence[54,55] and importance[56-58] of these lower abundance microbiota. As an example, *G. tenuis* is the type species of the recently described genus characterised within the human Gut Microbial Biobank constructed by Liu et al.[59] exploring "taxonomic dark matter" within the human gut microbiome. The authors predicted association of this previously uncultured species to weight-loss and highlighted it as worthy of further study in being low abundance but extremely widespread in humans (found in all datasets they investigated). The findings generated here, with common significant microbiome changes observed using 16 rRNA amplicon sequencing and WMS across a replicated biological question, reaffirm this importance of low abundance bacteria by suggesting these species could have roles in currently opaque pathologies, such as that induced by spaceflight.

#### 5 Intestinal cadherin gene expression during spaceflight

An exception to downregulation of cell adhesion molecules during spaceflight were cadherins, which included significant downregulation of *CLDN2* but up-regulation of *CLDN8*, *CLDN17* and *CLDN23*. Cadherins are transmembrane proteins concentrated in intestinal epithelia at tight junctions where they can form paracellular channels to mediate transport through intracellular spaces. Dysregulation of cadherins is associated with intestinal diseases such as IBD and disruption of epithelial integrity[110], and increases of the steroid hormone-regulated Claudin-8 (*CLDN8*) and Claudin-17 (*CLDN17*) have both been associated with tumorigenesis and cancer proliferation [111,112]. Interestingly, over-expression of *CLDN23* has recently been shown to improve intestinal barrier function but can decrease expression of *CLDN2*, notably involved in epithelial tight junction calcium absorption and bone mineralisation[113,114].

### 6 Supplementary figures

#### 6.1 Spaceflight alters murine gut microbiota

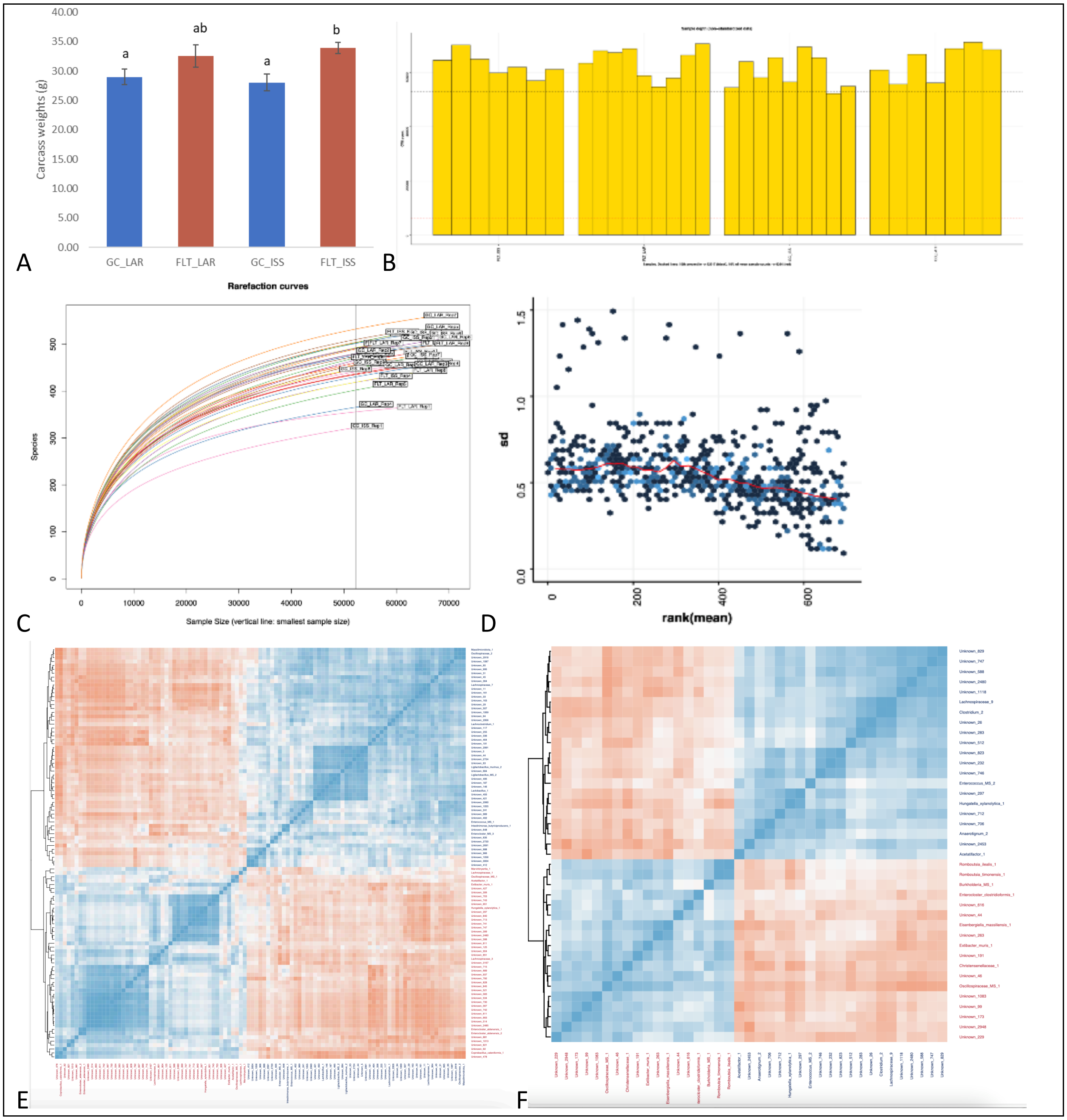

**Carcass weights and 16S rRNA gene amplification statistics** A) Carcass weights measured at dissection. B) 16S sample depth (i.e. ESV total abundance). Samples are grouped by replication. C) 16S rRNA rarefaction curves for each sample. D) Standard deviation of 16S normalized counts, i.e. the effect of rlog transformation on the variance. E) Pearson correlation of rlog transformed ESV abundance. Only differentially abundant ESVs are represented. The 2 colors represent the two different factors (GC LAR in blue and FLT LAR in red) F) Pearson correlation of rlog transformed ESV abundance. Only differentially abundant ESVs are represented. The 2 colors represent the two different factors (GC ISS in blue and FLT ISS in red).

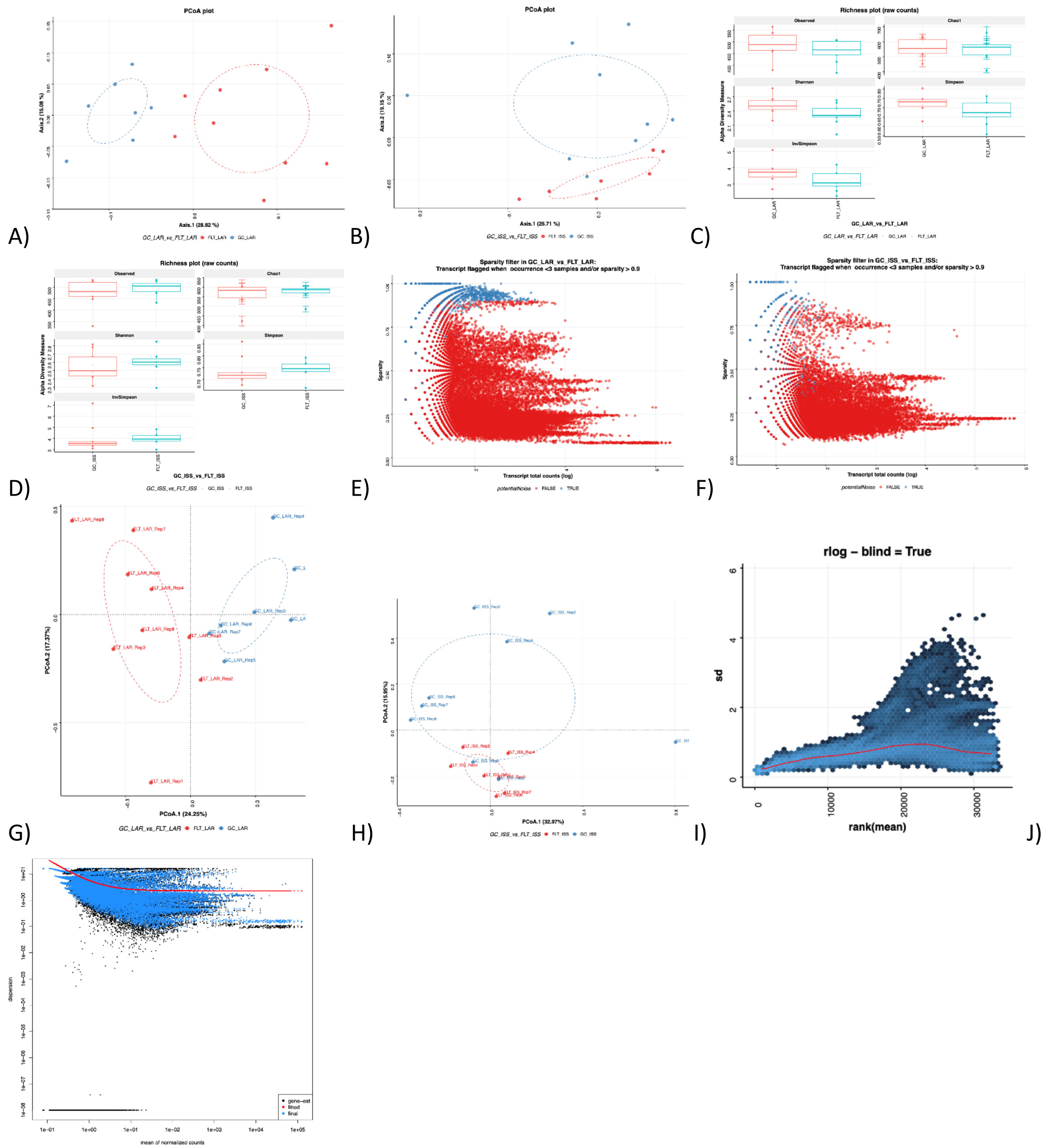

**16S rRNA gene amplification statistics (cont.)** A) Unsupervised ordination using rlog transformed ESV counts in LAR (16S rRNA). B) Unsupervised ordination using rlog transformed ESV counts in ISS (16S rRNA). C) 16S rRNA GC-LAR vs FLT-LAR alpha-diversity using multiple indexes. D) 16S rRNA GC-ISS vs FLT-ISS alpha-diversity using multiple indexes. E) Contig sparsity analysis in WGS LAR. Contig in blue were rejected from WGS statistical analysis. F) Contig sparsity analysis in WGS ISS. Contig in blue were rejected from WGS statistical analysis. G) Unsupervised ordination using cpm transformed contig counts in LARs (WGS). H) Unsupervised ordination using cpm transformed contig counts in ISS (WGS). I) Standard deviation of WGS normalized counts, i.e. the effect of cpm transformation on the variance. J) DESeq2 dispersion plot with the final estimates shrunk from the gene-wise estimates towards the fitted estimates.

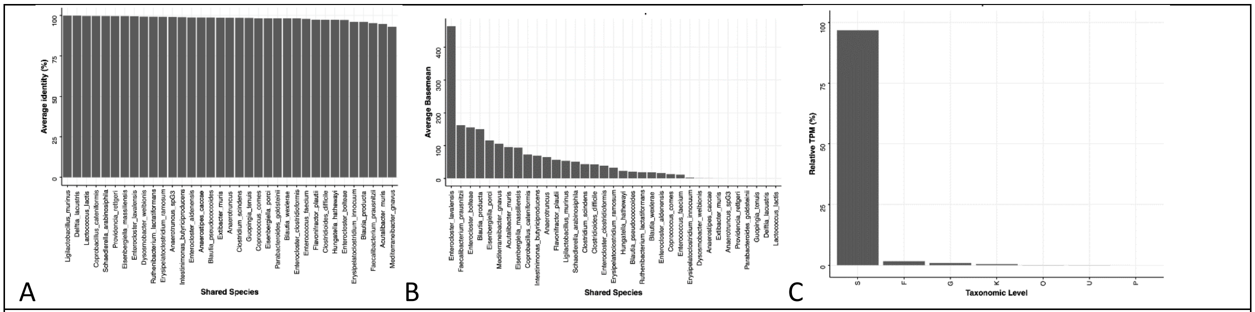


**WGS annotation statistics** A) WGS annotation. Average ANI per WGS species. B) Average normalized abundance (DESeq2 basemean) across WGS species. C) Relative CPM per taxonomic level

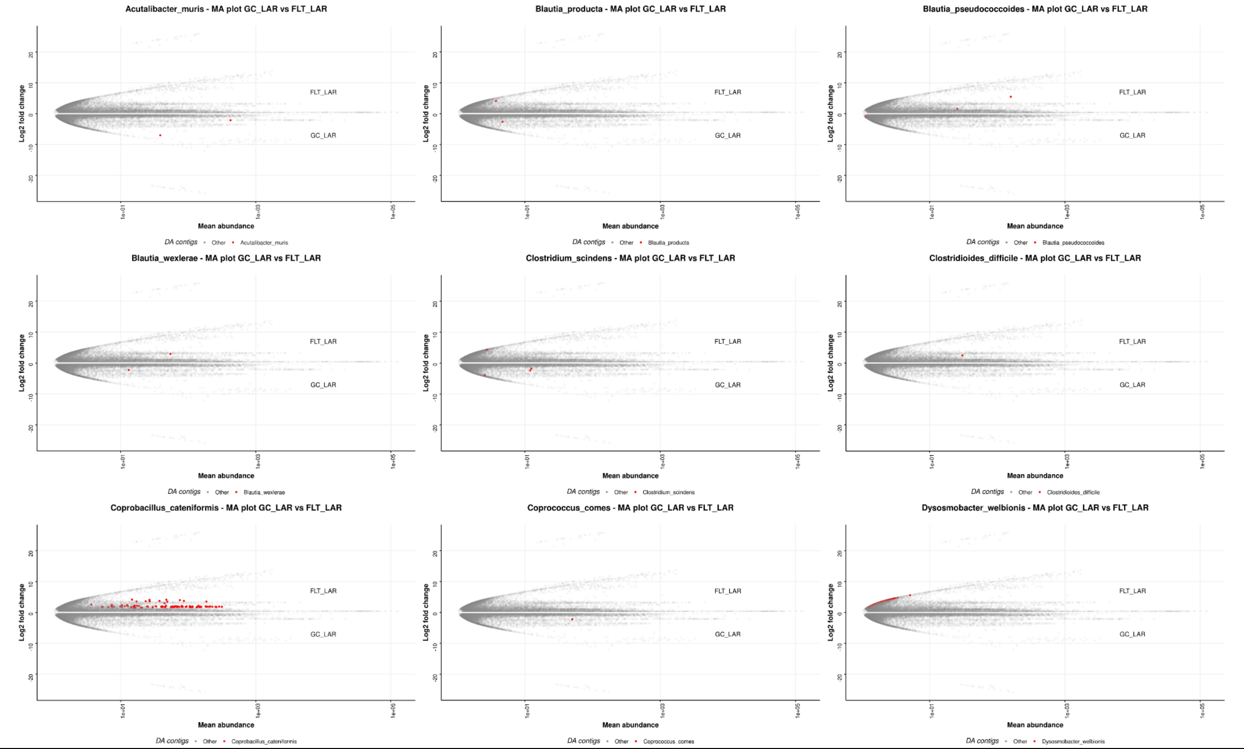

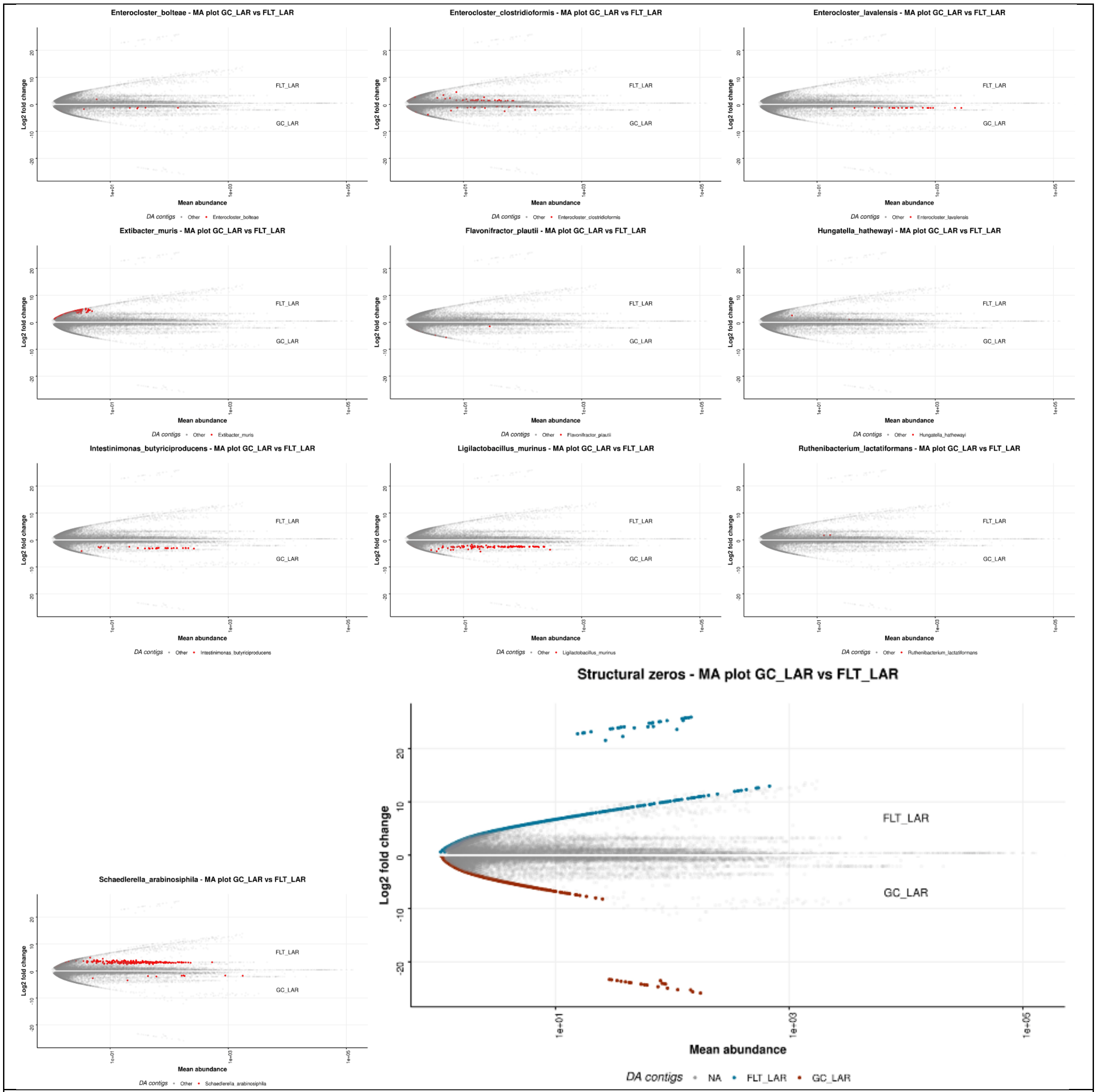


**MA plot of differentially abundant species in GC_LAR vs FLT_LAR differential abundance analysis (WGS)**

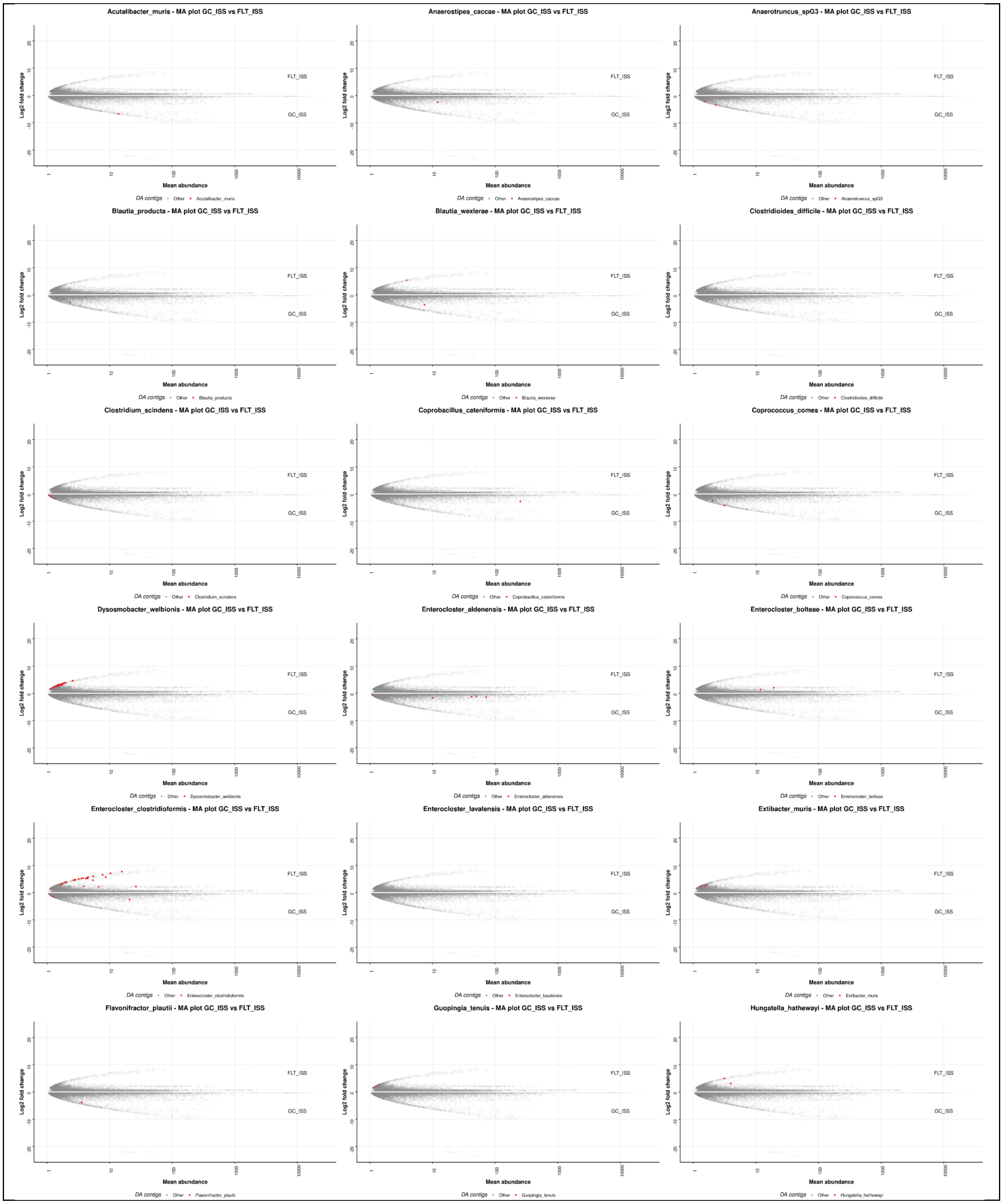
Acutalibacter_muris - MA plot GC_ISS vs FLT_ISS
Anaerostipes_caccae - MA plot GC_ISS vs FLT_ISS
Anaerotruncus_spG3 - MA plot GC_ISS vs FLT_ISS
Blautia_producta - MA plot GC_ISS vs FLT_ISS
Blautia_wexlerae - MA plot GC_ISS vs FLT_ISS
Clostridioides_difficile - MA plot GC_ISS vs FLT_ISS
Clostridium_scindens - MA plot GC_ISS vs FLT_ISS
Coprobacillus_cateniformis - MA plot GC_ISS vs FLT_ISS
Coprococcus_comes - MA plot GC_ISS vs FLT_ISS
Dysosmobacter_welbionis - MA plot GC_ISS vs FLT_ISS
Enterocloster_aldenensis - MA plot GC_ISS vs FLT_ISS
Enterocloster_bolteae - MA plot GC_ISS vs FLT_ISS
Enterocloster_clostridioformis - MA plot GC_ISS vs FLT_ISS
Enterocloster_lavalensis - MA plot GC_ISS vs FLT_ISS
Extibacter_muris - MA plot GC_ISS vs FLT_ISS
Flavonifractor_plautii - MA plot GC_ISS vs FLT_ISS
Guopingia_tenuis - MA plot GC_ISS vs FLT_ISS
Hungatella_hathewayi - MA plot GC_ISS vs FLT_ISS
FLT_ISS
GC_ISS
Log2 fold change
Mean abundance
DA contigs
Other

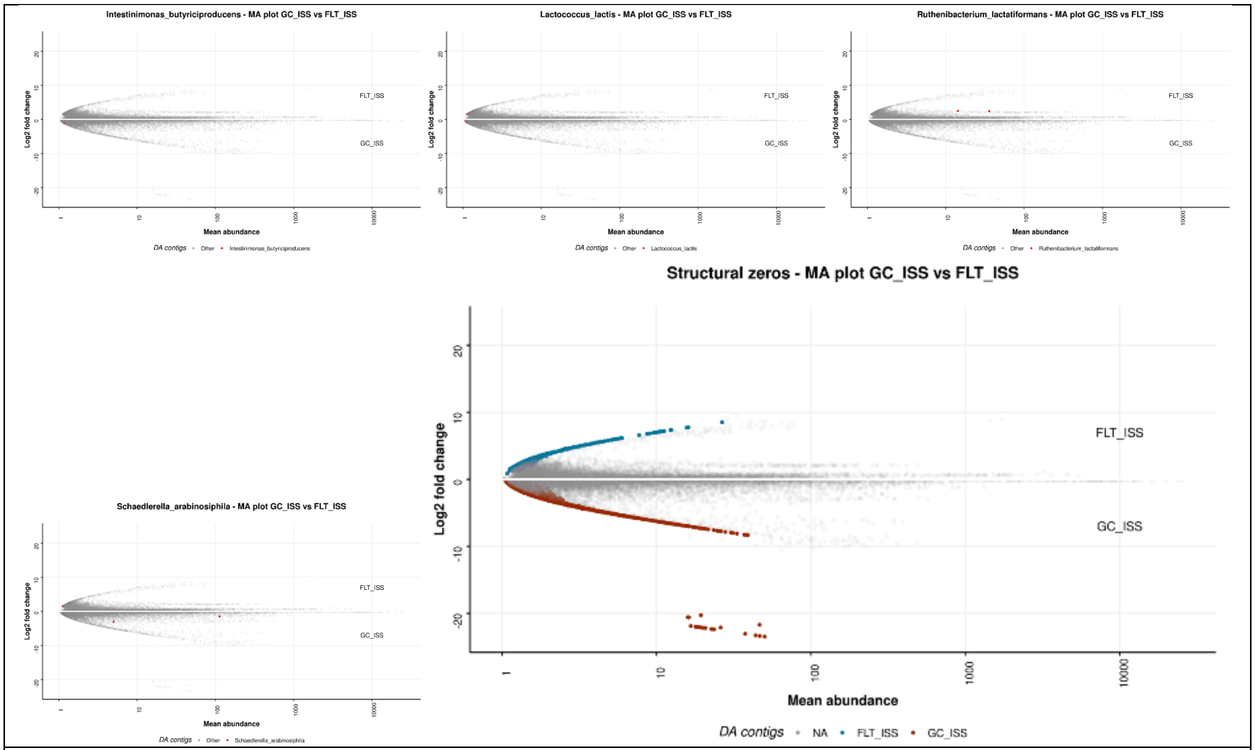


**MA plots of differentially abundant species in GC_ISS vs FLT_ISS differential abundance analysis (WGS)**

| **WGS functional pathways of interest** | |
|---|---|
| **KEGG Pathway** | Description |
| **52** | Galactose metabolism |
| **511** | Other glycan degradation |
| **531** | Glycosaminoglycan degradation |
| **520** | Amino sugar and nucleotide sugar metabolism |
| **500** | Starch and sucrose metabolism |
| **4512** | ECM-receptor interaction |
| **4350** | TGF-beta signaling pathway |
| **4976** | Bile secretion |
| **1212** | Fatty acid metabolism |
| **61** | Fatty acid biosynthesis |
| **51** | Fructose and mannose metabolism |
| **512** | Mucin type O-glycan biosynthesis |
| **511** | Other glycan degradation |
| **4973** | Carbohydrate digestion and absorption |
| **650** | Butanoate metabolism |
| **531** | Glycosaminoglycan degradation |
| **71** | Fatty acid degradation |
| **2025** | Biofilm formation - Pseudomonas aeruginosa |
| **121** | Secondary bile acid biosynthesis |

| | |
|---|---|
| **5100** | Bacterial invasion of epithelial cells |
| **1501** | beta-Lactam resistance |
| **1055** | Biosynthesis of vancomycin group antibiotics |
| **998** | Biosynthesis of various antibiotics |
| **4979** | Cholesterol metabolism |
| **515** | Mannose type O-glycan biosynthesis |
| **5130** | Pathogenic Escherichia coli infection |
| **ko02042** | Bacterial toxins |
| **ko01003** | Glycosyltransferases |
| **ko01504** | Antimicrobial resistance genes |
| **ko01011** | Peptidoglycan biosynthesis and degradation proteins |
| **ko00536** | Glycosaminoglycan binding proteins |
| **M00374** | Dicarboxylate-hydroxybutyrate cycle |
| **M00375** | Hydroxypropionate-hydroxybutylate cycle |
| **M00106** | Conjugated bile acid biosynthesis; cholate => taurocholate_glycocholate |
| **M00082** | Fatty acid biosynthesis; initiation |
| **M00083** | Fatty acid biosynthesis; elongation |
| **M00061** | D-Glucuronate degradation; D-glucuronate => pyruvate + D-glyceraldehyde 3P |
| **M00631** | D-Galacturonate degradation |
| **M00552** | D-galactonate degradation; De Ley-Doudoroff pathway; D-galactonate => glycerate-3P |
| **M00104** | Bile acid biosynthesis; cholesterol => cholate_chenodeoxycholate |
| **M00056** | O-glycan biosynthesis; mucin type core |
| **M00872** | O-glycan biosynthesis; mannose type |
| **M00019** | Valine_isoleucine biosynthesis; pyruvate => valine _ 2-oxobutanoate => isoleucine |
| **M00537** | Xylene degradation; xylene => methylbenzoate |

#### 6.2 Spaceflight alters host intestinal gene expression

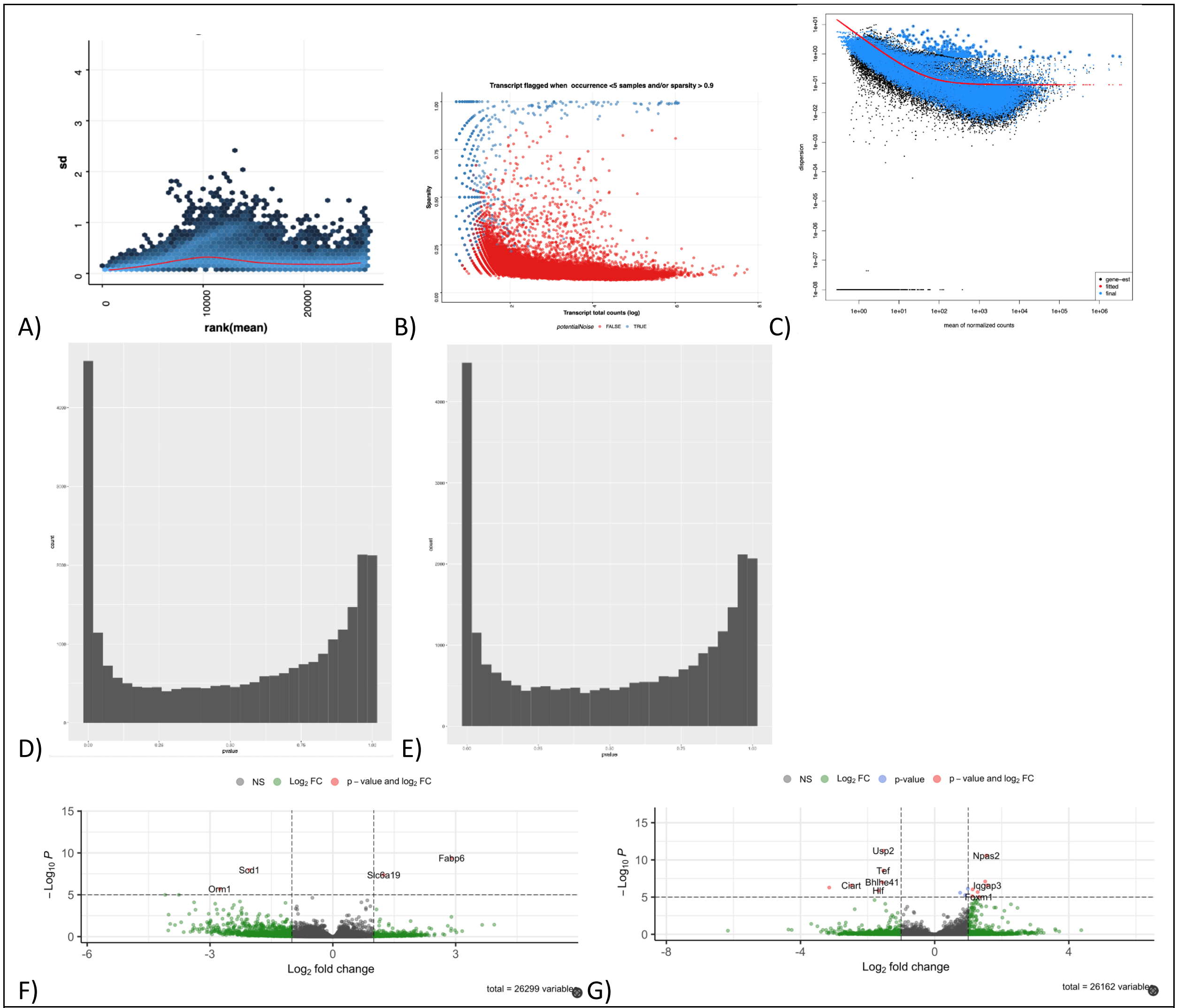


**Colon RNASeq statistics** A) Standard deviation of RNA-Seq normalized transcript counts (colon), i.e. the effect of rlog transformation on the variance. B) Sparsity analysis RNA-seq colon. Transcript in blue were rejected from statistical analysis. C) RNA-Seq DESeq2 dispersion plot (colon). D) DESeq2 P-value histogram in GC-LAR vs FLT-LAR (colon). E) DESeq2 P-value histogram in GC-ISS vs FLT-ISS (colon). F) Volcano plot of differential expression analysis of GC-LAR vs FLT-LAR. G) Volcano plot of differential expression analysis of GC-ISS vs FLT-ISS.

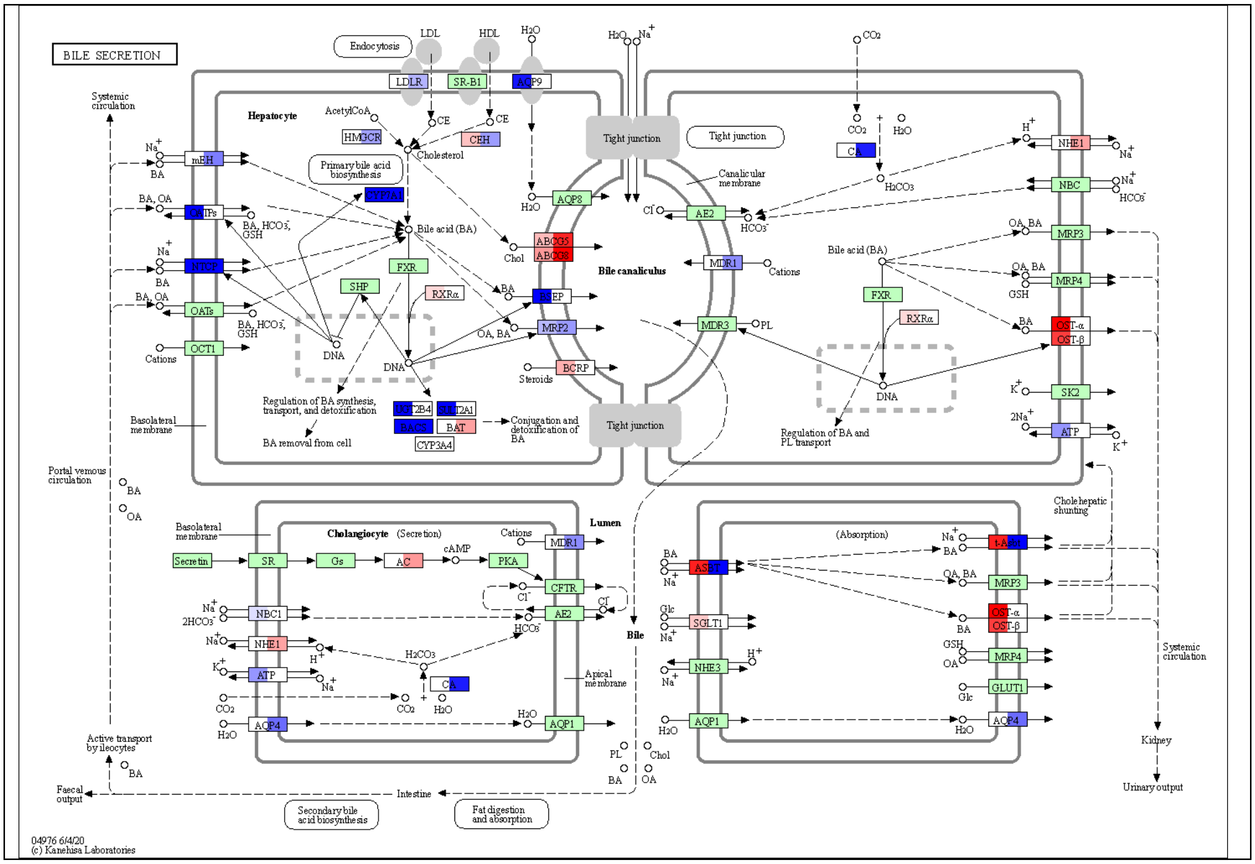
BILE SECRETION
Endocytosis
LDL
HDL
H2O
LDLR
SR-B1
AQP9
Systemic circulation
Hepatocyte
AcetylCoA
HMGCR
CE
CEH
Cholesterol
Primary bile acid biosynthesis
CYP7A1
mEH
OATPs
NTCP
OATs
OCT1
Na+
BA
BA, OA
BA, HCO3-, GSH
Cations
Bile acid (BA)
FXR
SHP
RXRα
DNA
Chol
ABCG5
ABCG8
BSEP
MRP2
OA, BA
BCRP
Steroids
Regulation of BA synthesis, transport, and detoxification
BA removal from cell
CYP2B4
SULT2A1
BAAT
BAT
CYP3A4
Conjugation and detoxification of BA
Basolateral membrane
Tight junction
AQP8
Cl-
AE2
HCO3-
MDR1
MDR3
PL
Bile canaliculus
Canalicular membrane
CO2
H2O
CA
H2CO3
H+
NHE1
NBC
HCO3-
MRP3
MRP4
OST-α
OST-β
SK2
ATP
K+
2Na+
GSH
Regulation of BA and PL transport
Portal venous circulation
OA
Cholangiocyte
(Secretion)
Secretin
SR
Gs
AC
cAMP
PKA
CFTR
NBC1
2HCO3-
NHE1
ATP
AQP4
AQP1
Lumen
Apical membrane
Bile
(Absorption)
t-ASBT
ASBT
SGLT1
Glc
NHE3
GLUT1
Cholehepatic shunting
Systemic circulation
Kidney
Urinary output
Active transport by ileocytes
Faecal output
Intestine
Secondary bile acid biosynthesis
Fat digestion and absorption
04976 6/4/20
(c) Kanehisa Laboratories

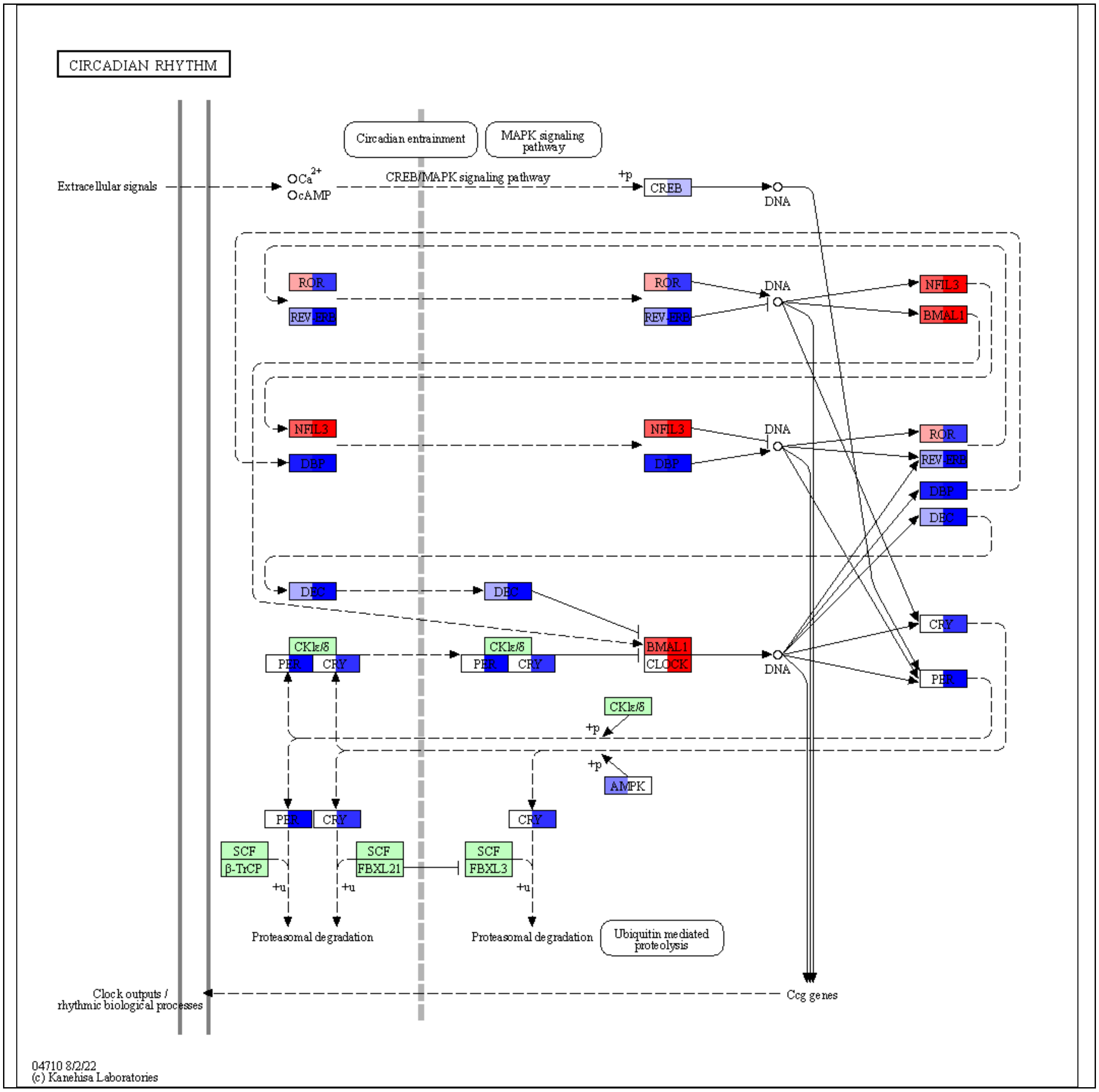
CIRCADIAN RHYTHM
Circadian entrainment
MAPK signaling pathway
Extracellular signals
Ca2+
cAMP
CREB/MAPK signaling pathway
+p
CREB
DNA
ROR
REV-ERB
NFIL3
BMAL1
DBP
DEC
CRY
PER
CKIε/δ
CLOCK
AMPK
SCF
β-TrCP
FBXL21
FBXL3
+u
Proteasomal degradation
Ubiquitin mediated proteolysis
Clock outputs / rhythmic biological processes
Ccg genes
04710 8/2/22
(c) Kanehisa Laboratories

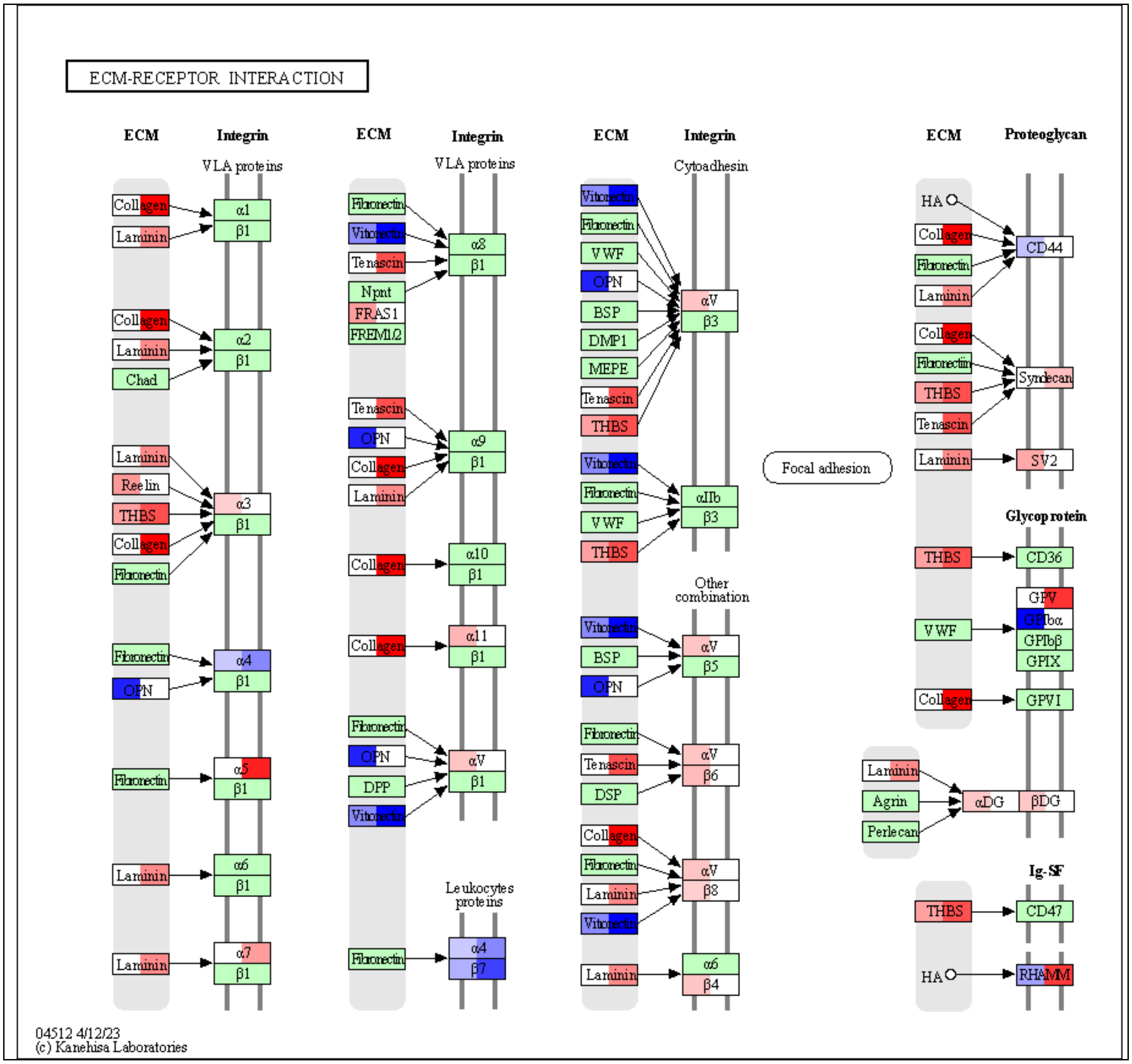

ECM-RECEPTOR INTERACTION
ECM
Integrin
VLA proteins
Cytoadhesin
Other combination
Leukocytes proteins
Proteoglycan
Glycoprotein
Ig-SF
Focal adhesion
Collagen
Laminin
Chad
Reelin
THBS
Fibronectin
OPN
Vitronectin
Tenascin
Npnt
FRAS1
FREM1/2
DPP
VWF
BSP
DMP1
MEPE
DSP
HA
Agrin
Perlecan
CD44
Syndecan
SV2
CD36
GPV
GPIbα
GPIbβ
GPIX
GPVI
αDG
βDG
CD47
RHAMM
α1
α2
α3
α4
α5
α6
α7
α8
α9
α10
α11
αV
αIIb
β1
β3
β4
β5
β6
β7
β8
04512 4/12/23
(c) Kanehisa Laboratories

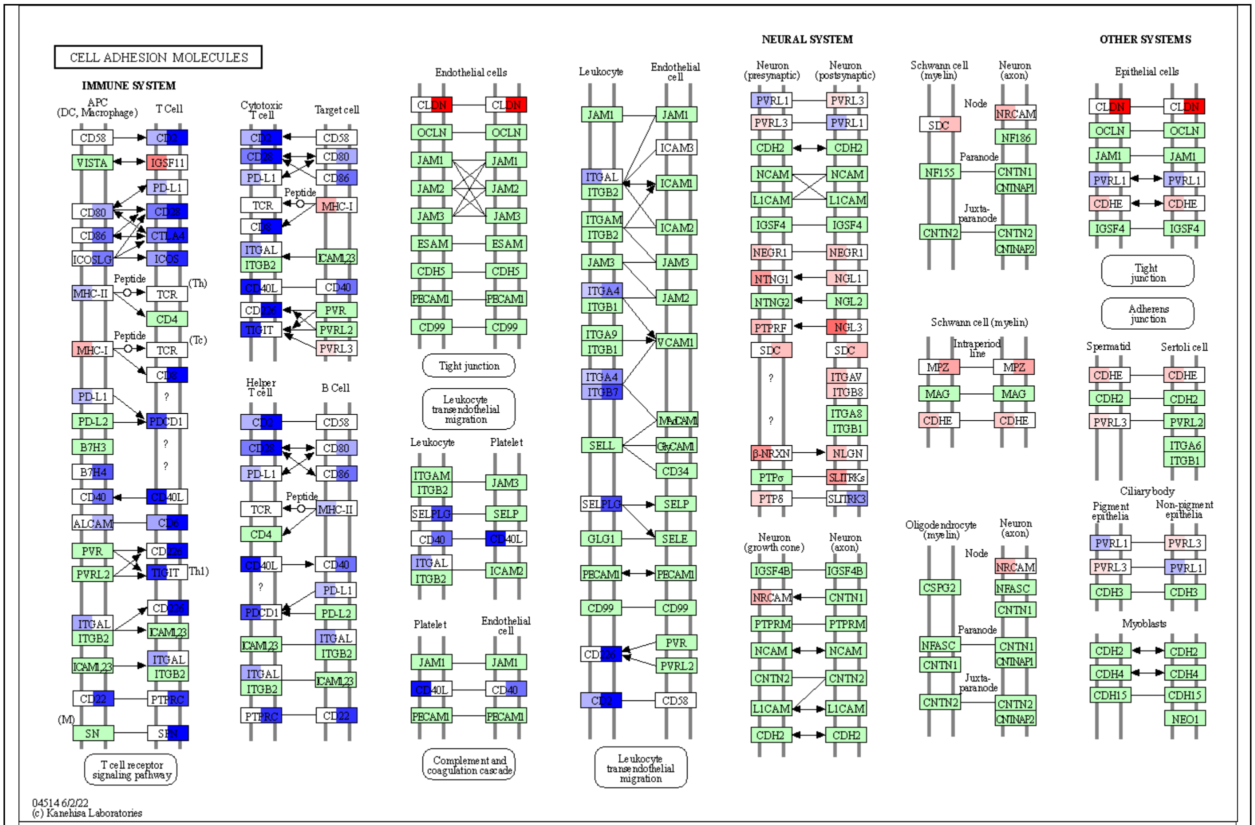
CELL ADHESION MOLECULES
IMMUNE SYSTEM
NEURAL SYSTEM
OTHER SYSTEMS
APC (DC, Macrophage)
T Cell
Cytotoxic T cell
Target cell
Helper T cell
B Cell
Endothelial cells
Leukocyte
Endothelial cell
Platelet
Neuron (presynaptic)
Neuron (postsynaptic)
Schwann cell (myelin)
Neuron (axon)
Epithelial cells
Tight junction
Leukocyte transendothelial migration
Complement and coagulation cascade
T cell receptor signaling pathway
Adherens junction
Spermatid
Sertoli cell
Ciliary body
Pigment epithelia
Non-pigment epithelia
Myoblasts
Oligodendrocyte (myelin)
Neuron (growth cone)
04514 6/2/22
(c) Kanehisa Laboratories

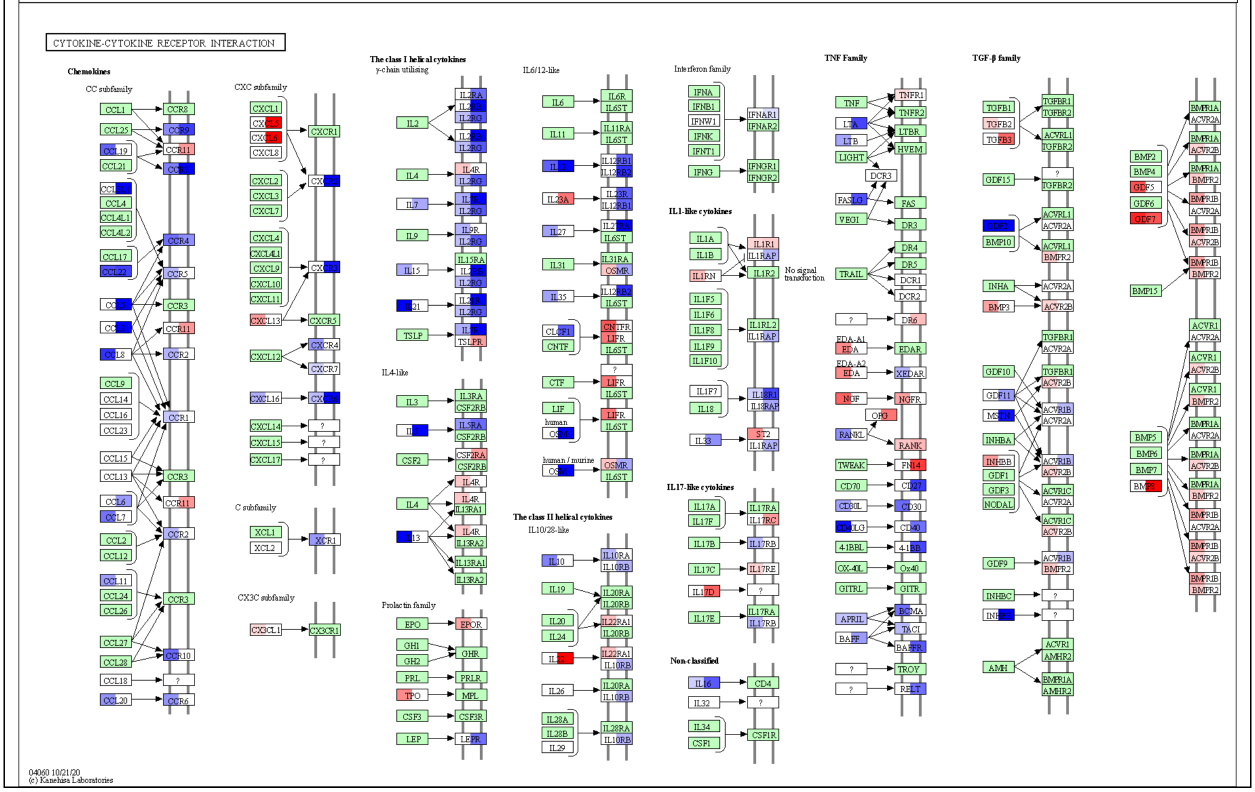
CYTOKINE-CYTOKINE RECEPTOR INTERACTION
Chemokines
CC subfamily
CXC subfamily
C subfamily
CX3C subfamily
The class I helical cytokines
γ-chain utilizing
IL4-like
Prolactin family
IL6/12-like
The class II helical cytokines
IL10/28-like
Interferon family
IL1-like cytokines
IL17-like cytokines
Non-classified
TNF Family
TGF-β family
04060 10/21/20
(c) Kanehisa Laboratories

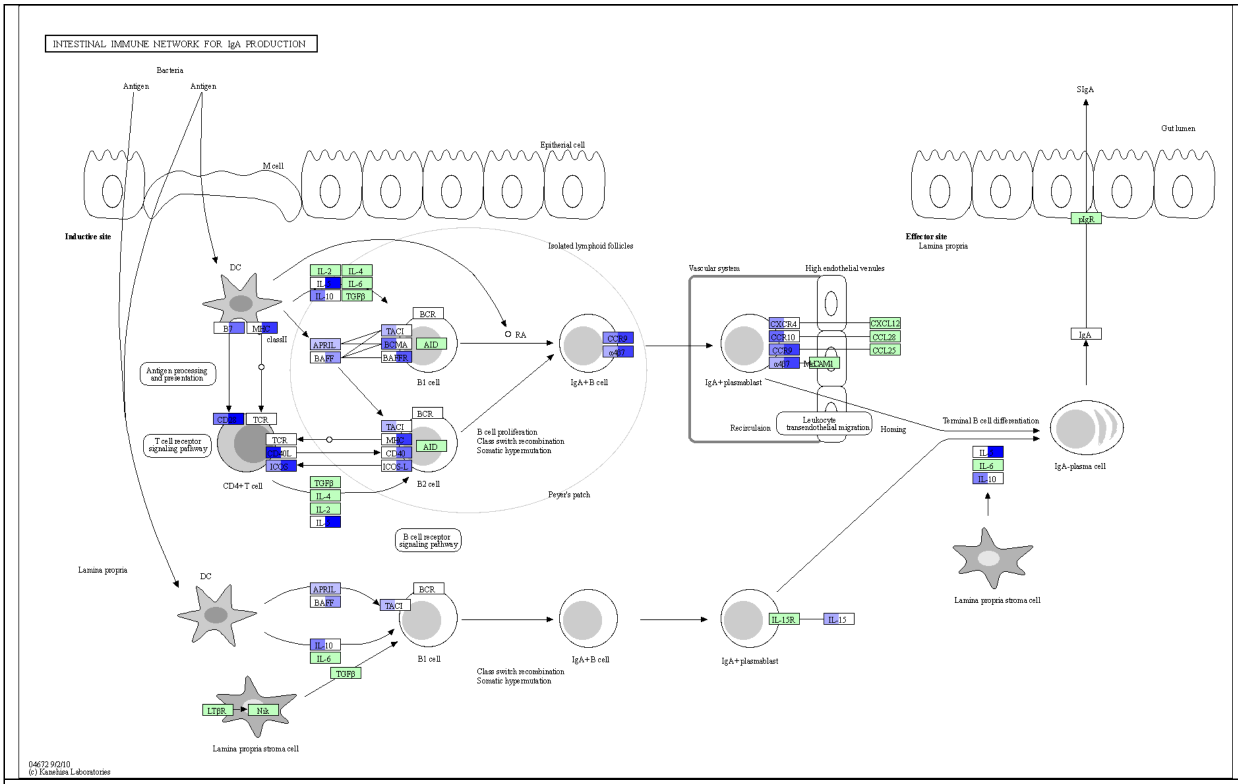


**Selected KEGG pathways with colon differentially expressed genes**. Blue is downregulated and red is upregulated (padj<0.1); left half of gene is GC_LAR vs FLT_LAR (29 days of spaceflight) and right half of gene is GC_ISS vs FLT_ISS (56 days of spaceflight). See *RR6 host transcriptomics* file for full DE gene list.

#### 6.3 Spaceflight alters gene expression in the liver

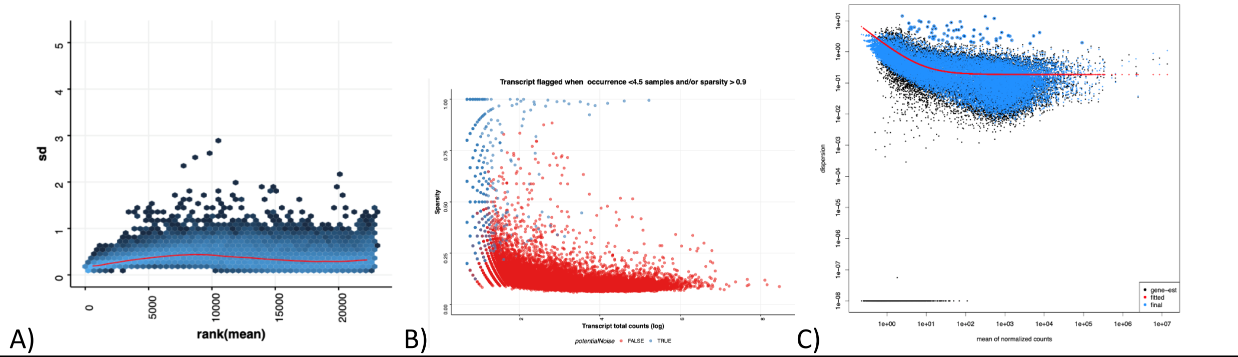

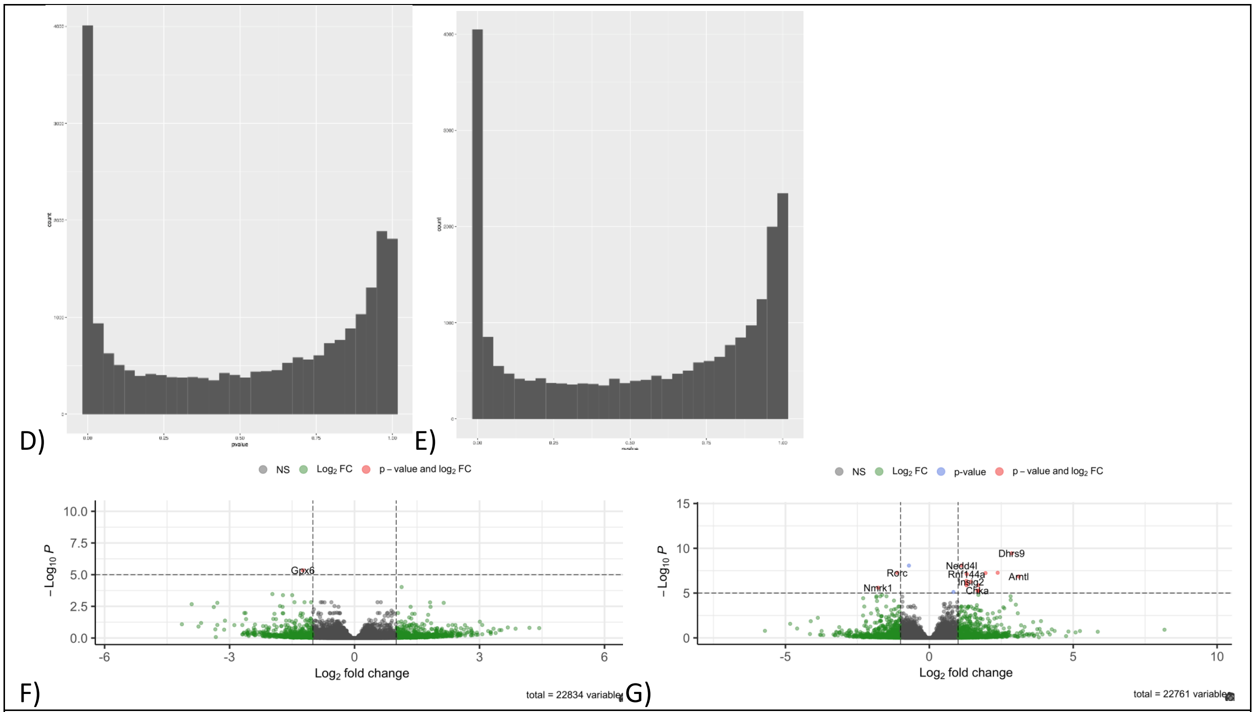


**Liver RNASeq statistics** A) Standard deviation of RNA-Seq normalized transcript counts (liver), i.e. the effect of rlog transformation on the variance. B) Sparsity analysis RNA-seq liver. Transcript in blue were rejected from statistical analysis. C) RNA-Seq DESeq2 dispersion plot (liver). D) DESeq2 P-value histogram in GC-LAR vs FLT-LAR (liver). E) DESeq2 P-value histogram in GC-ISS vs FLT-ISS (liver). F) Volcano plot of differential expression analysis of GC-LAR vs FLT-LAR. G) Volcano plot of differential expression analysis of GC-ISS vs FLT-ISS.

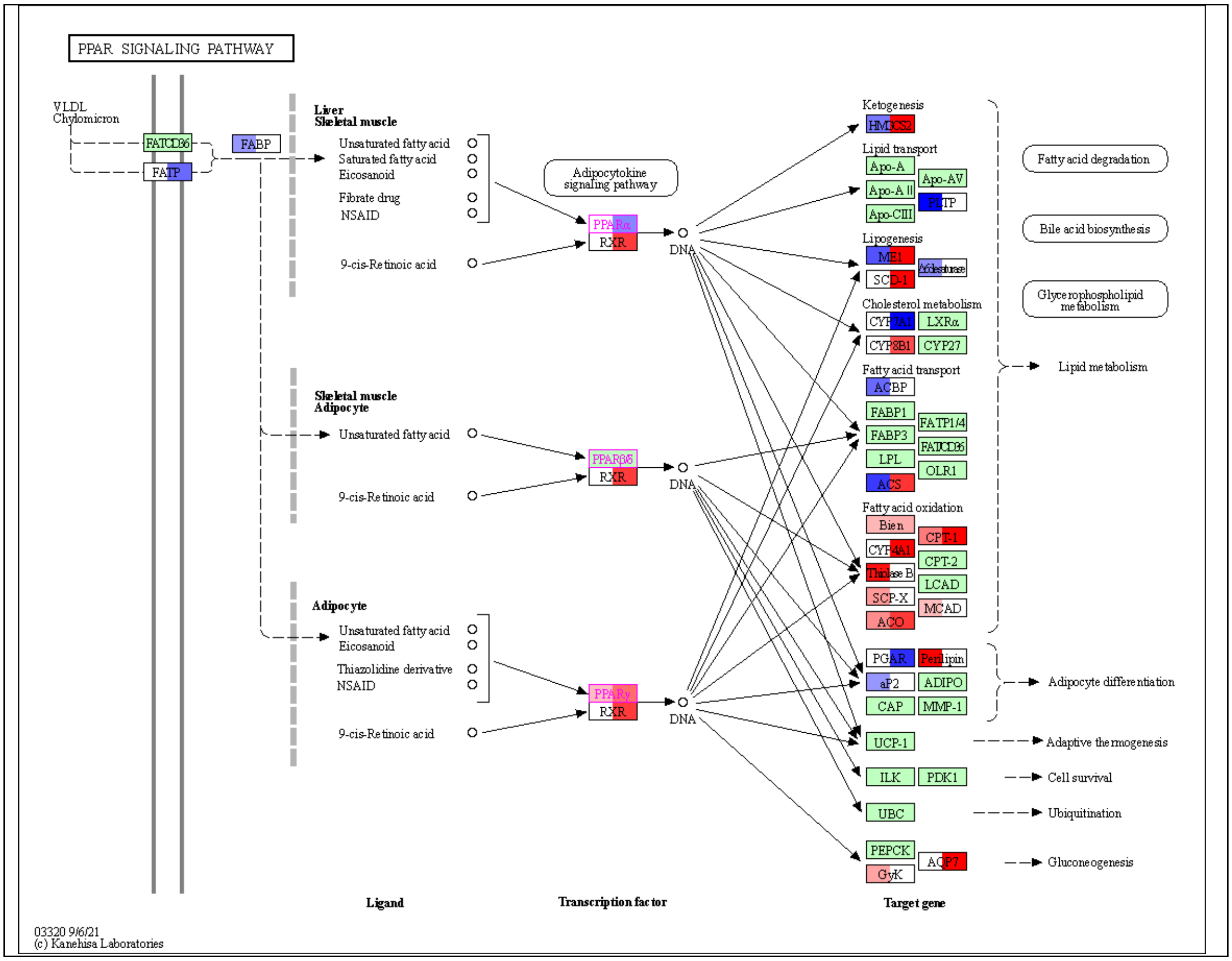
PPAR SIGNALING PATHWAY
VLDL
Chylomicron
FAT/CD36
FATP
FABP
Liver
Skeletal muscle
Unsaturated fatty acid
Saturated fatty acid
Eicosanoid
Fibrate drug
NSAID
9-cis-Retinoic acid
Adipocytokine signaling pathway
PPARα
RXR
DNA
Skeletal muscle
Adipocyte
Unsaturated fatty acid
9-cis-Retinoic acid
PPARβ/δ
RXR
DNA
Adipocyte
Unsaturated fatty acid
Eicosanoid
Thiazolidine derivative
NSAID
9-cis-Retinoic acid
PPARγ
RXR
DNA
Ketogenesis
HMGCS2
Lipid transport
Apo-A
Apo-AV
Apo-A II
PLTP
Apo-CIII
Lipogenesis
ME1
Δ6desaturase
SCD-1
Cholesterol metabolism
CYP7A1
LXRα
CYP8B1
CYP27
Fatty acid transport
ACBP
FABP1
FATP1/4
FABP3
FAT/CD36
LPL
OLR1
ACS
Fatty acid oxidation
Bien
CPT-1
CYP4A1
CPT-2
Thiolase B
LCAD
SCP-X
MCAD
ACO
PGAR
Perilipin
aP2
ADIPO
CAP
MMP-1
UCP-1
ILK
PDK1
UBC
PEPCK
AQP7
GyK
Fatty acid degradation
Bile acid biosynthesis
Glycerophospholipid metabolism
Lipid metabolism
Adipocyte differentiation
Adaptive thermogenesis
Cell survival
Ubiquitination
Gluconeogenesis
Ligand
Transcription factor
Target gene
03320 9/6/21
(c) Kanehisa Laboratories

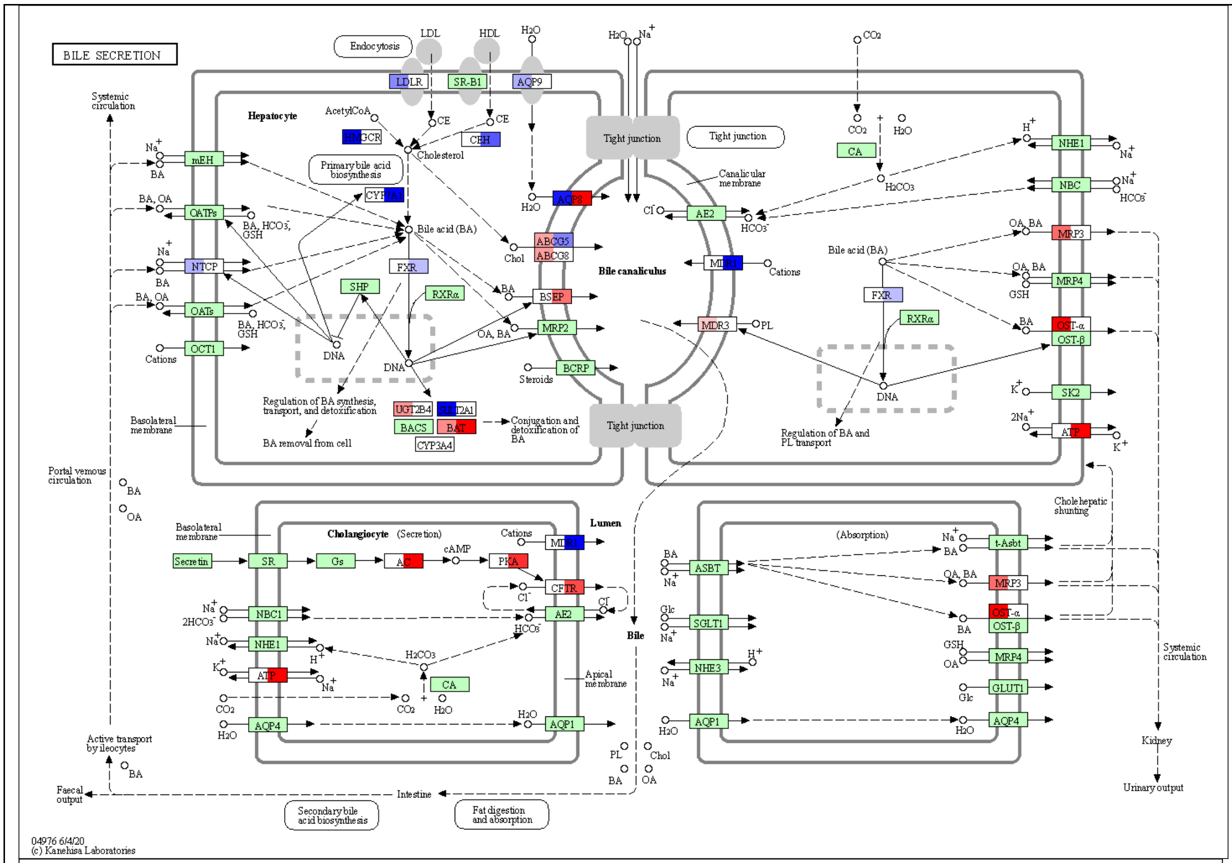
BILE SECRETION
Endocytosis
LDL
HDL
LDLR
SR-B1
AQP9
Systemic circulation
Hepatocyte
AcetylCoA
CE
CEH
Cholesterol
Primary bile acid biosynthesis
Bile acid (BA)
Tight junction
Canalicular membrane
Bile canaliculus
mEH
OATPs
NTCP
OATs
OCT1
FXR
SHP
RXRα
DNA
Cations
Chol
ABCG5
ABCG8
BSEP
MRP2
BCRP
MDR3
AE2
CA
NHE1
NBC
MRP3
MRP4
OST-α
OST-β
SK2
ATP
Steroids
Regulation of BA synthesis, transport, and detoxification
UGT2B4
BACS
BAT
CYP3A4
Conjugation and detoxification of BA
BA removal from cell
Basolateral membrane
Regulation of BA and PL transport
Portal venous circulation
Cholangiocyte
(Secretion)
Secretin
SR
Gs
AC
cAMP
PKA
CFTR
NBC1
NHE1
AQP4
AQP1
Apical membrane
Lumen
Bile
(Absorption)
ASBT
SGLT1
NHE3
t-Asbt
GLUT1
Cholehepatic shunting
Kidney
Urinary output
Active transport by ileocytes
Faecal output
Intestine
Secondary bile acid biosynthesis
Fat digestion and absorption
04976 6/4/20
(c) Kanehisa Laboratories

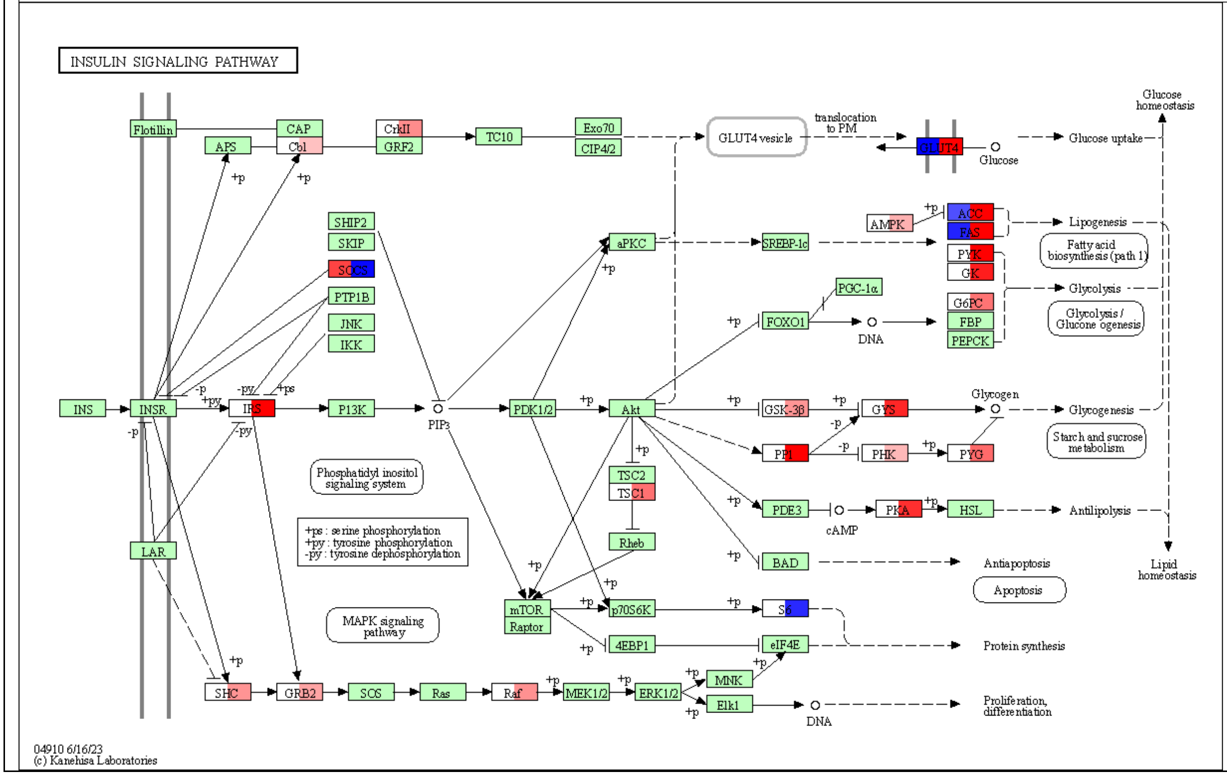
INSULIN SIGNALING PATHWAY
Flotillin
APS
CAP
Cbl
CrkII
GRF2
TC10
Exo70
CIP4/2
GLUT4 vesicle
translocation to PM
GLUT4
Glucose
Glucose uptake
Glucose homeostasis
SHIP2
SKIP
SOCS
PTP1B
JNK
IKK
aPKC
SREBP-1c
AMPK
ACC
FAS
PYK
GK
Lipogenesis
Fatty acid biosynthesis (path 1)
Glycolysis
PGC-1α
FOXO1
G6PC
FBP
PEPCK
Glycolysis / Gluconeogenesis
INS
INSR
IRS
PI3K
PIP3
PDK1/2
Akt
GSK-3β
GYS
Glycogen
Glycogenesis
PP1
PHK
PYG
Starch and sucrose metabolism
TSC2
TSC1
Rheb
PDE3
cAMP
PKA
HSL
Antilipolysis
Lipid homeostasis
BAD
Antiapoptosis
Apoptosis
Phosphatidyl inositol signaling system
+ps : serine phosphorylation
+py : tyrosine phosphorylation
-py : tyrosine dephosphorylation
LAR
mTOR
Raptor
p70S6K
S6
4EBP1
eIF4E
Protein synthesis
MAPK signaling pathway
SHC
GRB2
SOS
Ras
Raf
MEK1/2
ERK1/2
MNK
Elk1
DNA
Proliferation, differentiation
04910 6/16/23
(c) Kanehisa Laboratories

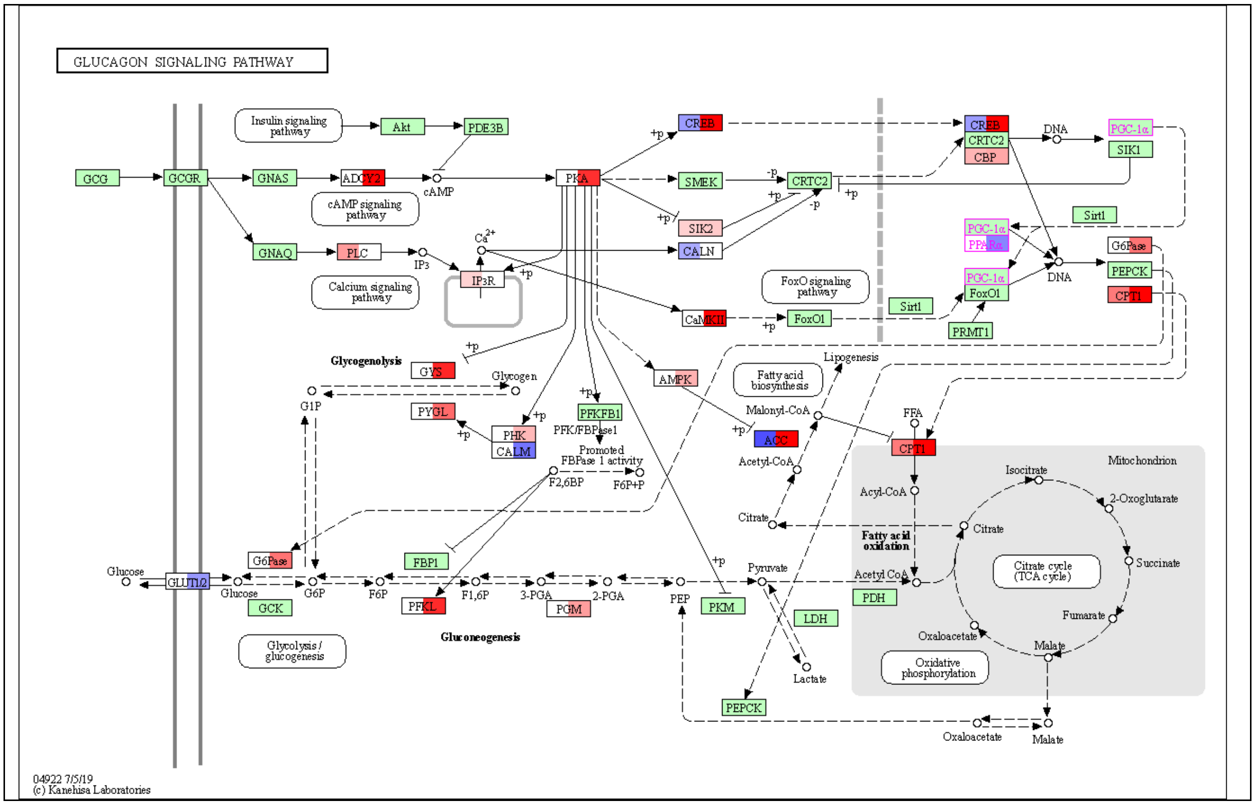
GLUCAGON SIGNALING PATHWAY
Insulin signaling pathway
Akt
PDE3B
GCG
GCGR
GNAS
ADCY2
cAMP
PKA
CREB
SMEK
CRTC2
SIK2
CALN
cAMP signaling pathway
GNAQ
PLC
IP3
Ca2+
IP3R
CaMKII
FoxO1
Calcium signaling pathway
FoxO signaling pathway
CREB
CRTC2
CBP
DNA
PGC-1α
SIK1
Sirt1
PGC-1α
PPARα
PGC-1α
FoxO1
PRMT1
Sirt1
DNA
G6Pase
PEPCK
CPT1
Glycogenolysis
GYS
Glycogen
G1P
PYGL
PHK
CALM
PFKFB1
PFK/FBPase1
Promoted FBPase 1 activity
F2,6BP
F6P+P
AMPK
Fatty acid biosynthesis
Lipogenesis
Malonyl-CoA
ACC
Acetyl-CoA
Citrate
FFA
CPT1
Mitochondrion
Isocitrate
2-Oxoglutarate
Acyl-CoA
Citrate
Fatty acid oxidation
Succinate
Citrate cycle (TCA cycle)
Acetyl CoA
PDH
Fumarate
Oxaloacetate
Malate
Oxidative phosphorylation
Glucose
GLUT2
Glucose
G6Pase
G6P
F6P
FBP1
F1,6P
3-PGA
2-PGA
PEP
Pyruvate
GCK
PFKL
PGM
PKM
LDH
Lactate
Glycolysis / gluconeogenesis
Gluconeogenesis
PEPCK
Oxaloacetate
Malate
04922 7/5/19
(c) Kanehisa Laboratories

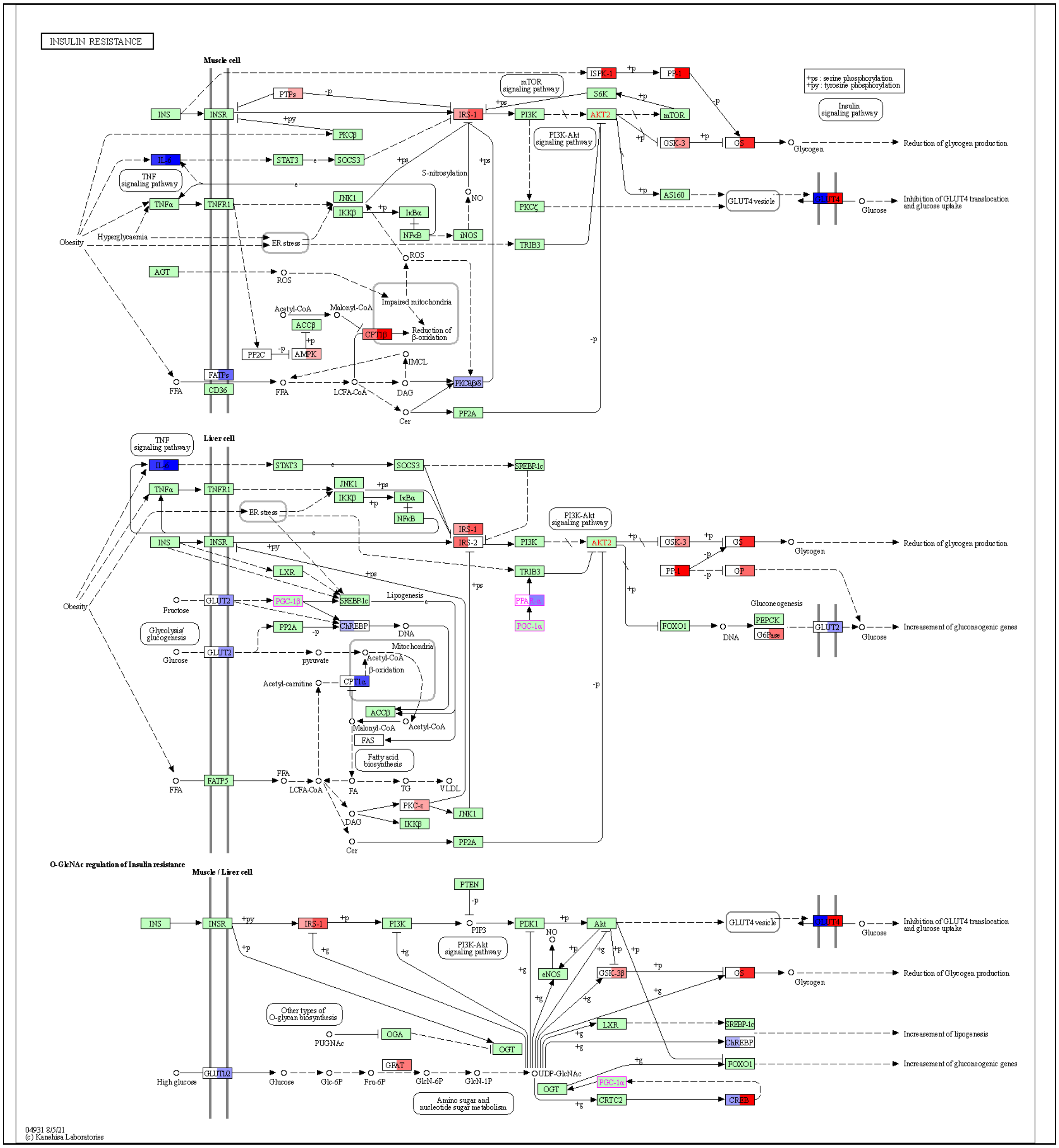
INSULIN RESISTANCE
Muscle cell
+ps : serine phosphorylation
+py : tyrosine phosphorylation
Insulin signaling pathway
mTOR signaling pathway
PI3K-Akt signaling pathway
TNF signaling pathway
INS
INSR
PTPs
PKCβ
STAT3
SOCS3
TNFα
TNFR1
JNK1
IKKβ
IκBα
NFκB
iNOS
S-nitrosylation
NO
IRS-1
PI3K
AKT2
S6K
mTOR
GSK-3
AS160
PKCζ
TRIB3
GLUT4 vesicle
Glycogen
Glucose
Reduction of glycogen production
Inhibition of GLUT4 translocation and glucose uptake
Obesity
Hyperglycaemia
ER stress
AGT
ROS
Acetyl-CoA
Malonyl-CoA
ACCβ
AMPK
PP2C
Impaired mitochondria
Reduction of β-oxidation
IMCL
FFA
FATPs
CD36
LCFA-CoA
DAG
Cer
PP2A
Liver cell
SREBP-1c
IRS-2
LXR
GLUT2
Fructose
Glycolysis/ glucogenesis
PGC-1β
ChREBP
DNA
Lipogenesis
Mitochondria
β-oxidation
pyruvate
Acetyl-carnitine
FAS
Fatty acid biosynthesis
FATP5
FA
TG
VLDL
PKC-ε
PGC-1α
FOXO1
Gluconeogenesis
PEPCK
Increasement of gluconeogenic genes
O-GlcNAc regulation of Insulin resistance
Muscle / Liver cell
PTEN
PIP3
PDK1
Akt
eNOS
GSK-3β
Other types of O-glycan biosynthesis
OGA
PUGNAc
OGT
High glucose
GLUT1/2
Glc-6P
Fru-6P
GFAT
GlcN-6P
GlcN-1P
UDP-GlcNAc
Amino sugar and nucleotide sugar metabolism
CRTC2
Reduction of Glycogen production
Increasement of lipogenesis
04931 8/5/21
(c) Kanehisa Laboratories

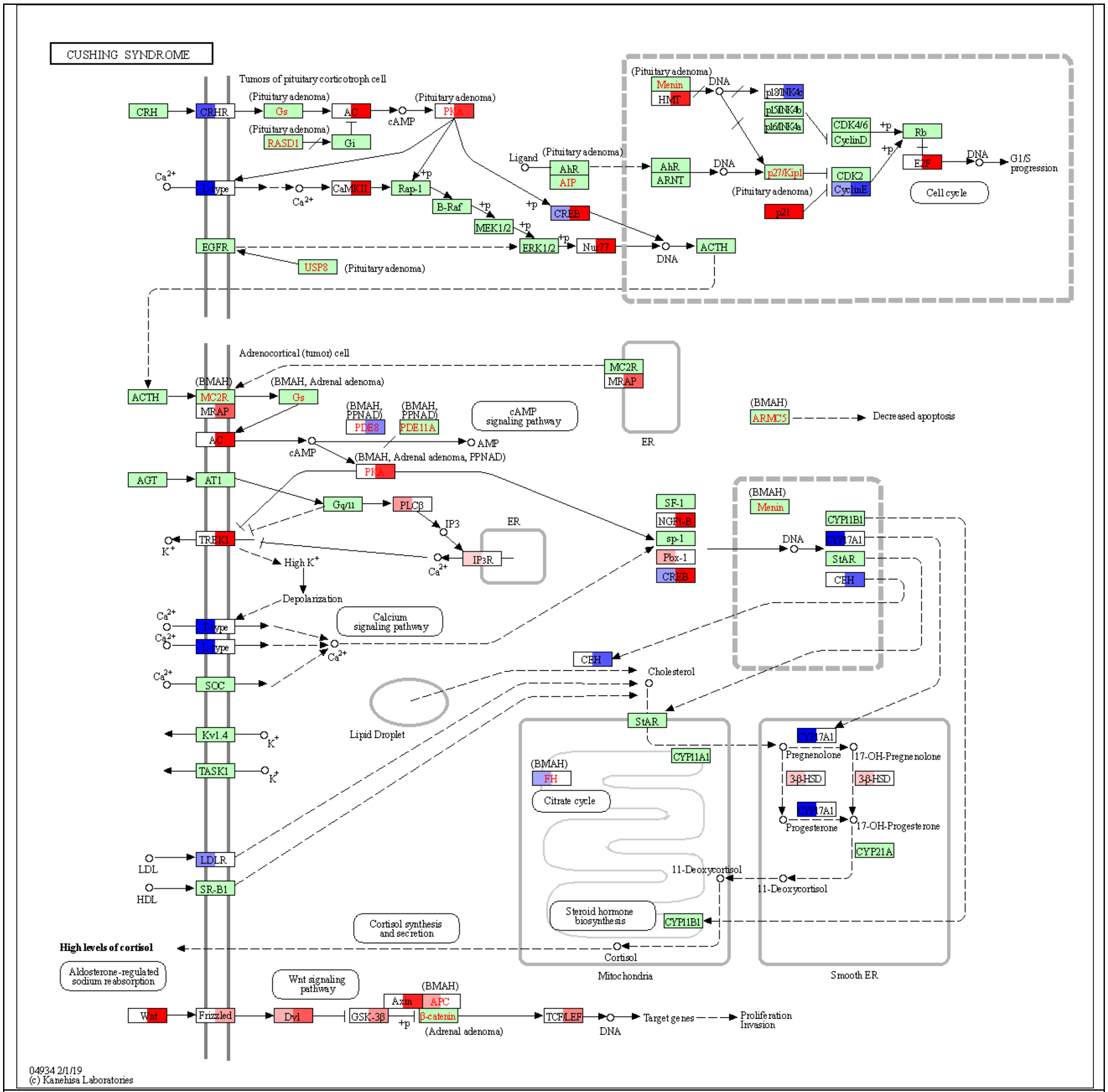


**Selected KEGG pathways with liver differentially expressed genes**. Blue is downregulated and red is upregulated (padj<0.1); left half of gene is GC_LAR vs FLT_LAR (29 days of spaceflight) and right half of gene is GC_ISS vs FLT_ISS (56 days of spaceflight). See *RR6 host transcriptomics* file for full DE gene list.
